# Effects of social presence on behavioural, neural and physiological aspects of empathy for pain

**DOI:** 10.1101/2022.09.14.507943

**Authors:** Pauline Petereit, Ronja Weiblen, Anat Perry, Ulrike M. Krämer

## Abstract

In mediated interactions (e.g. video calls), less information is available about the other. To investigate how this affects our empathy for one another, we conducted an EEG study, in which thirty human participants observed one of 5 targets undergoing painful electric stimulation, once in a direct interaction and once in a live, video-mediated interaction. We found that observers were as accurate in judging others’ pain and showed as much affective empathy via video as in a direct encounter. While mu suppression, a common neural marker of empathy, was not sensitive to others’ pain, theta responses to others’ pain as well as skin conductance coupling between participants were reduced in the video-mediated condition. We conclude that physical proximity with its rich social cues is important for nuanced physiological resonance with the other’s experience. More studies are warranted to confirm these results and to understand their behavioural significance for remote social interactions.

## Introduction

Over the last decades, many social interactions in private life and at work, including medical and psychotherapeutic contexts, have shifted from personal encounters to mediated interactions. In contrast to a direct interaction, a mediated interaction is not happening face-to-face in the same space, but instead mediated via any means of communication, for example a letter, a phone call, or a video call. These interactions differ in two aspects from personal encounters: First, they provide less detailed social cues and information channels, a feature often termed intimacy. Second, they limit opportunities for immediate, reciprocal exchange compared to personal encounters, a feature often termed immediacy. These factors contribute together to the degree of social presence offered within social interactions. Note that focusing on them extends the concept of social presence from the dichotomous presence or absence of another person to a feature of the medium that can vary continuously (Short et al. 1976; Cui et al. 2013; but see Biocca et al. 2003 for alternative accounts of social presence). Given the reduced social presence of mediated interactions, the question arises of how they change our ability to share and understand others’ affective states, i.e., our ability to empathize (Decety and Jackson 2004; Shamay-Tsoory 2011). Focusing on intimacy, we asked how the reduction of social cues – using a video call versus a direct, personal interaction – affects empathy. While visual and auditory cues are usually still present in a video mediated interaction, visual cues are limited as people can only see a cutting of the scene and not the whole body and environment of the other. Moreover, the resolution and other quality aspects of visual and auditory data might be reduced in a video call. In addition, olfactory and tactile information, which are thought to be important for empathy, are completely missing in a video call (Goldstein et al. 2018; Calvi et al. 2020). To investigate the effects of changed social presence on behavioural, neural and physiological aspects of empathy, we used empathy for pain as a well-established model (Singer and Lamm 2009).

Empathy for pain is a multifaceted process, including an affective response, feelings of distress and empathic care towards the person suffering (Lamm et al. 2007; Goubert et al. 2009; Singer and Lamm 2009), as well as cognitive processes. The latter are sometimes measured as empathic accuracy, which is the accuracy of one’s perception of the other’s pain (J. Zaki et al. 2009; Laursen et al. 2014).

On the neural level, empathy for pain has been hypothesized to be supported by representing the other’s bodily state in one’s own somatosensory system and thus with neural activations in that system while observing others in pain (Riečanský and Lamm 2019). These neural activations are associated with mu suppression in the EEG: reduced power between 8 and 13 Hz over the somatosensory cortex (Cheng et al. 2008; Perry, Bentin, et al. 2010; Peled-Avron et al. 2018; Peng et al. 2021). Mu suppression has been linked to higher empathic accuracy (Goldstein et al. 2018). Although one study has shown that activity in the somatosensory cortex is sensitive to others’ painful facial expressions (Gallo et al. 2018), most studies have used static pictures or videos of injured body limbs (Whitmarsh et al. 2011; Motoyama et al. 2017; Levy et al. 2018; Zebarjadi et al. 2021) and it is therefore not clear whether this phenomenon is specific to the perception of others’ injured bodies. It is therefore useful to additionally examine neural activity at other frequencies or areas of the cortex that might be linked to pain empathy. Some studies examined midfrontal theta activity (4–8 Hz) as a possible electrophysiological shared component of empathy (Mu et al., 2008; Peng et al., 2021) and own pain experience (Misra et al. 2017; Ploner et al. 2017; Peng et al. 2021). Mid-frontal theta is thought to indicate activity in the anterior cingulate cortex (ACC, Mitchell et al. 2008; van der Molen et al. 2017), which is often reported in fMRI studies on empathy for pain and is associated with the negative affect during own and others’ pain (Fallon et al. 2020). ACC activation is also sensitive to facial expressions of different pain intensities (Saarela et al. 2007).

Finally, several studies suggest that physiological “coupling” (in cardiac activity or skin conductance), i.e., aligning to the physiological state of someone in pain, might reflect empathic sharing and facilitate understanding of the other (Goldstein et al. 2017; Jospe et al. 2020; Reddan et al. 2020 Apr 17; Zerwas et al. 2021).

Humans share others’ emotional state especially in response to social cues such as facial expressions or eye contact (Ensenberg et al., 2017; Hess, 2021; Perry et al., 2010b). We thus expected that reduced availability of social cues and thereby reduced intimacy of the interaction would dampen affective responses and impair understanding of others. Additionally, if people do not share the same space, with the potential of physical interaction, feelings of closeness and the salience of the other person and with that also empathic responses might be diminished (Melloni et al. 2014; Järvelä et al. 2016; Weisz and Zaki 2018; Bogdanova et al. 2022). To test the effect of intimacy on empathy for pain in the current study, pairs of participants underwent an empathy-for-pain paradigm in two conditions: one direct, face-to-face interaction and one interaction mediated via real-time video transfer. In both conditions, one participant (the target) received painful electrical stimulation to the back of the right hand while the other (the observer) was watching. As both observer and target were able to see each other in both conditions, their behaviours could subtly influence each other. Therefore, the task can be considered an interactive empathy task (Cui et al. 2013; Cañigueral et al. 2022 Jun 1). The task of the observer was to judge the pain intensity experienced by the target and her/his own feelings of unpleasantness while watching the others’ pain. As targets themselves also rated their experienced pain intensity, the correspondence between targets’ and observers pain provided a behavioural read-out for empathic accuracy for pain. In addition, we measured EEG from the observer and cardiac and skin conductance responses from both members of the dyad.

## Methods

### Participants

Five female psychology students were recruited as targets. Only females were sampled for the targets to reduce possible gender effects over the dyads. They were on average 19.8 years old (SD = 0.75). Thirty psychology students (7 males, 23 females, mean age(SD) = 24.07(4.68) years) were recruited as observers. As all the students were from the same university and most of them from the same year, some of them had already met before the experiment (n =10), while others had not (n =20). A priori power analyses with data simulations using the *simr* package (Green and MacLeod 2016) in R showed that this sample size is sufficient to detect a small effect (*f*_2_ = 0.02) of condition on empathic accuracy (measured via an interaction effect between shock intensity and direct vs. mediated condition in a generalized linear mixed model on single trials; see below for statistical analyses) with a power of at least 0.98 (depending on exact model structure). Data were simulated using sample parameters (means and variance parameters as well as correlations between conditions) from a pilot study. We did not conduct a separate power analysis for the neural data because effect size estimations for neural effects are difficult. Moreover, the neural analysis was conducted with the full amount of trials (80 trials) in contrast to the 40 trials used for the behavioural analysis. Hence, the power calculation for the behavioural analyses is more conservative. For both targets and observers, exclusion criteria were current psychiatric or cardiovascular and past or current neurological disorders, current or chronic pain conditions or current pain-medication intake. For one observer, all physiological data from the own-pain condition had to be excluded because of missing stimulus triggers. Skin conductance and electrocardiogram (ECG) data from one target (mediated-interaction condition) were missing due to a technical error. Skin conductance data from two observers could not be analysed due to poor data quality. These dyads were excluded from all analyses of the missing outcome variable. Targets received €10 per hour; observers received course credit or €10 per hour. All participants provided written informed consent prior to taking part in the study, including consent for video-recording them and showing the videos to other work group members, and in case of the targets, showing the videos to other participants in future studies. The experiment was carried out according to the Declaration of Helsinki and was approved by the Ethics Committee of the University of Lübeck.

### Experimental design

Each target interacted with six different observers on six different study days. Observers came to the lab once for one session (two interactions) with one target. For the observers, there were three conditions, which were implemented in separate blocks (each observer went through all three conditions, within-subject design, see Fig. 1A). In the “own pain” condition, the observer was alone in the laboratory and received electric shocks. This procedure ensured that observers knew what the electric shocks felt like. In the “direct interaction” condition, observer and target sat opposite each other at a table, and the target received electric shocks while the observer watched. Thus, in this condition the observers were able to see the whole body of the target apart from the legs, which were placed under the table. In the “mediated interaction” condition, target and observer sat in adjacent rooms and saw each other over a real-time video transmission. The video showed the face, upper body, arms and hands of the target, and targets were instructed to place the hand with the shock electrode clearly visible in front of them on the table in both conditions. As the camera filming the target had to be placed above the screen the target was watching, the visual perspective on the target was slightly different between the “mediated” and the “direct” condition. In both conditions, a blue board was placed behind the target to make the background as similar as possible. Observers watched the same target in the latter two conditions and rated the observed pain experience of the target. Targets rated their own pain experience. In all conditions, skin conductance and ECG were recorded from both participants, and EEG was recorded from the observer. The “own pain” condition was always carried out first, and the order of “direct interaction” and “mediated interaction” conditions was pseudo-randomized over participants such that each order (“direct interaction” first, or “mediated interaction” first) occurred an equal number of times. In the latter two conditions, targets’ and observers’ behaviour was video-recorded.

**Figure 1.**
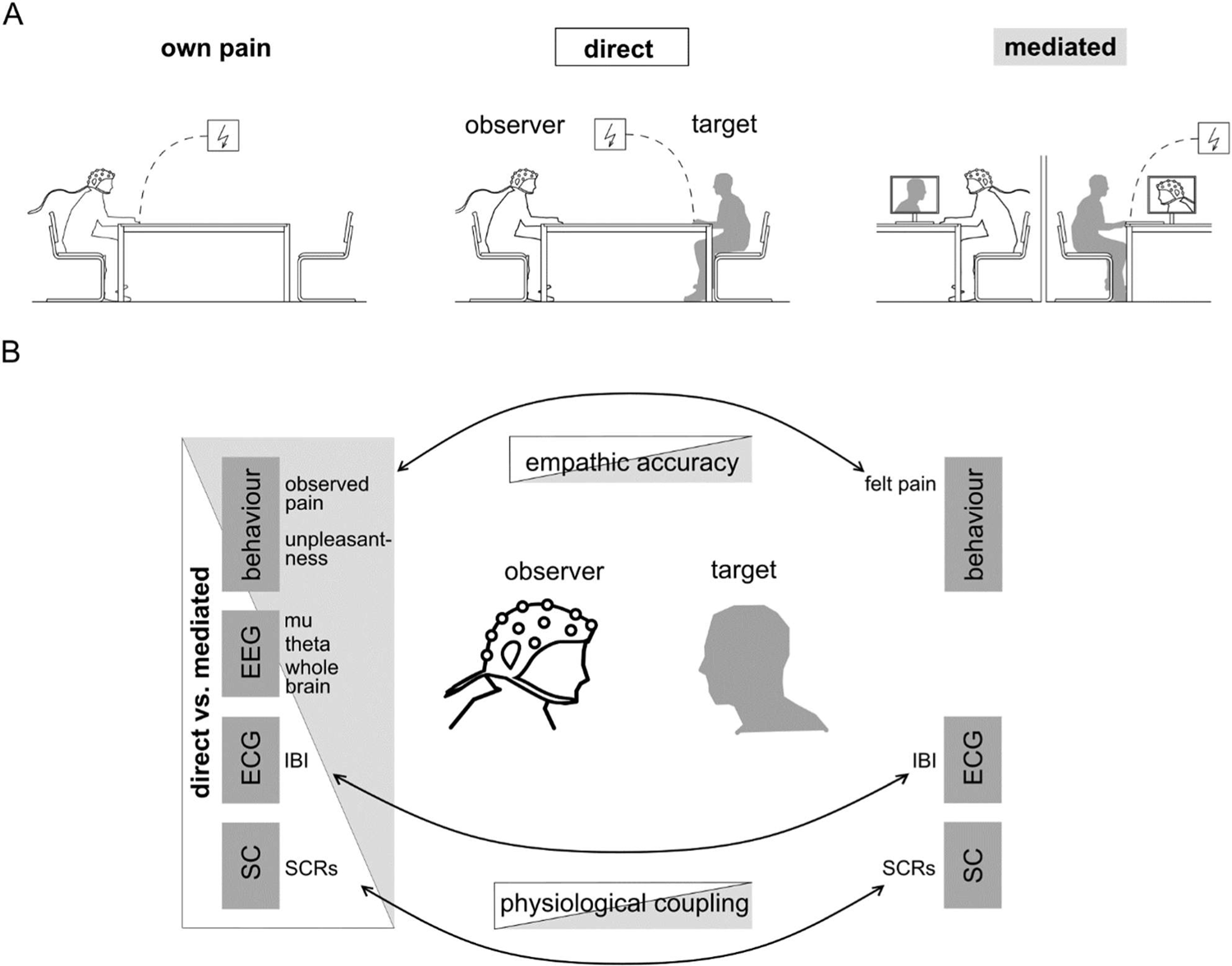
(A) Schematic overview of experimental conditions. (B) Overview of the analysed outcomes.

### Stimulus calibration

Prior to the pain task, pain stimuli were calibrated to the subjective pain thresholds of the participants (the observers in the “own pain” condition, and the targets prior to the first interaction condition). To this end, participants received electric shocks starting from 0 mA, increasing in amplitude in steps of 0.5 mA. They were required to rate each stimulus on a 9-point Likert scale from 0 (“not perceivable”) to 8 (“strongest pain imaginable”). The intermediate steps on the scale were likewise labelled (1 = “noticeable”, 2 = “unpleasant”, 3 = “slightly painful”, 4 = “medium painful”, 5 = “very painful”, 6 = “extremely painful”, 7 = “unbearably painful”). We increased the stimulus intensity until participants rated a stimulus with “7” (Rutgen et al. 2015). After that, stimuli were decreased in amplitude (again in steps of 0.5 mA) until participants rated the stimulus as “0” or an amplitude of 0 mA was reached. The procedure was then repeated once more with increasing stimulus intensity. The stimulus intensity that was rated as 1 (“noticeable”) in this last round was used as the lower limit for the stimuli presented during the task, with the stimulus intensity rated as 6 (“extremely painful”) used as the upper limit. For the pain task itself, the range from the lower limit to the upper limit was evenly divided into 20 levels, to have a finer variation in shock intensity levels. As the physical intensity of the stimuli was increased until participants rated them as “extremely painful”, we could make sure that participants really experienced pain during the experiment.

### Pain task

The pain task itself was adapted from Rutgen and colleagues (2015). Electric shocks were delivered using a DS 5 isolated bipolar constant current stimulator (Digitimer) and a bar electrode (Digitimer, two electrodes with 9-mm diameter, 30 mm apart) attached to the back of the right hand. The skin under the electrode was treated with an abrasive paste and conductive gel to reduce the electric resistance of the skin.

Each trial started with an auditory cue lasting 500 ms that did not predict the shock stimulus intensity. At 1000 ms after the cue, the electric shock was delivered for 500 ms (series of 2-ms electric pulses, interspersed with approximately 20-ms breaks). After a randomly varying interval (6000–9000 ms), the next trial or the rating followed. In 50% of the trials, participants were prompted to rate the stimulus by a vocal recording saying, “Please rate”. We included the rating only in 50% of the trials to keep the duration of the experiment feasible. The rating was given on a tablet computer. Targets rated how painful the electric shock was for themselves on a visual analogue scale ranging from “not at all painful” to “extremely painful”. Observers rated how painful they thought the electric shock was for the target (on the same visual analogue scale) as well as how unpleasant it was for them to watch the target receive the electric shock (on a visual analogue scale ranging from “not at all unpleasant” to “extremely unpleasant”). The latter rating served as a measure of affective empathy. Electric shocks varied in intensity in 20 steps from the intensity the targets had rated as “noticeable” (intensity level 1) to the intensity they had rated as “extremely painful” (intensity level 20) during the calibration. There were 80 trials in each condition, and each intensity occurred four times. The number of trials was similar to other EEG studies on pain empathy with a similar paradigm (Rutgen et al. 2015). However, compared to former studies, we enhanced the number of rated trials from one third to half of the trials, because our power analysis had shown that this was necessary for a reliable assessment of empathic accuracy. The order of intensities was pseudo-random (with no more than four shocks with intensity level higher than 10 or lower than 11 in a row) but fixed for all participants and conditions. The order of trials that had to be rated was fixed as well. In the “own pain” condition, the task was the same except that observers rated their own pain experience on the visual analogue scale. Targets and observers were instructed not to talk or move excessively during the task but were otherwise allowed to express their emotions freely. Observers were instructed to rate the pain of the other as accurately as possible. We assumed that it would be possible for the observer to infer the intensity of targets’ pain from facial expressions, gestures, para-lingual expression and body movement. Observers were free to use all these cues as they wished. Both observers and targets were told who would receive shocks in the task, hence the observers knew that they would receive shocks in the “own pain” condition, but would not receive any shocks in the other two conditions.

### Experimental procedure

#### Target selection

Before targets interacted with observers, they came to the laboratory alone to familiarize themselves with the procedures and the stimuli. In this first session, they did the same pain task as in the main experiment, but with no other person in the room. As in the main experiment, skin conductance and ECG were recorded. After the first session, targets decided whether they wanted to participate in the main experiment. Moreover, we used the first session for target selection, as we aimed to recruit only targets who set the intensity limits of the stimuli during stimulus calibration to a level that was actually painful. This constraint was important to ensure that we measured actual pain empathy during the main experiment. We therefore defined the minimum upper intensity limit that targets had to reach during the first session to be eligible for the full experiment as within +/-1 standard deviation of the mean upper intensity limit of a pilot study (resulting in a minimum upper intensity limit of 2.5 mA). We invited seven potential targets to the first session. Due to the criteria, we had to exclude one target, and one participant dropped out after the first session, which left five targets for the main experiment.

#### Main experiment

For the main experiment, the observer arrived first in the laboratory. After the informed-consent form was signed, the EEG, skin conductance and ECG measurement equipment as well as the stimulus electrode were prepared. The pain-calibration procedure was carried out. After five practice trials to familiarize themselves with the ratings on the tablet computer, participants did the pain task in the “own pain” condition. Meanwhile, the target arrived in a different room, responded to questionnaires and was equipped with the electrodes for physiological measurement. As soon as the “own pain” condition was finished, the stimulus electrode was attached to the target’s right hand, and the target underwent the pain-calibration procedure. Meanwhile, the observer completed questionnaires. When both were finished, the experimental tasks started (see “pain task” above; either “direct interaction” or “mediated interaction” first). Between conditions, there was a short break in which the target moved between rooms (either into our out of the room in which the observer was seated). ECG and skin conductance cables of the observer were unplugged from the amplifier during moving, and plugged in again before the next condition started. Data quality was checked again before the second condition started. Afterwards, target and observer were seated in different rooms again and replied to post-experimental questionnaires. Finally, observers were debriefed about the aim of the study. Targets were debriefed only after completing all six sessions.

### Questionnaires

We assessed participants’ age, gender, body weight and height, educational degree, and habits regarding smoking, caffeine consumption and physical activity. After the experiment, we obtained participants’ subjective evaluation of the experiment and observers’ evaluation of the target. They also filled out two personality questionnaires: the Interpersonal Reactivity Index (Davis, 1983; German version: Paulus, 2009) and the Emotion Regulation Questionnaire (Gross & John, 2003; German version: Abler & Kessler, 2009). These data are not further evaluated here.

### Physiological data acquisition

Participants were asked to refrain from smoking, exercise, alcohol and caffeine for at least six hours before the experiment to prevent these factors from impacting the physiological measurements. EEG data were recorded with 59 Ag/AgCl electrodes placed on an elastic cap according to the international 10-20-system (using a BrainAmp MR plus amplifier, BrainProducts GmbH,). An online reference electrode was placed on the left earlobe, while an offline reference electrode was placed on the right earlobe. Horizontal and vertical EOG were recorded with four electrodes placed next to the outer corners of the eyes and above and below the left eye, respectively. Sampling rate was 500 Hz, and data were recorded with an online high pass filter of 0.016 Hz, a low pass filter of 48 Hz and a notch filter at 50 Hz. Impedances were kept below 5kΩ.

ECG data were recorded with bipolar Ag/AgCl recording electrodes and one reference electrode, using a 50-Hz notch filter. One of the recording electrodes was placed on the right forearm, the other one on the left lower calf of the participant (following Einthoven lead II configuration, Einthoven et al., 1950). Skin conductance was measured with two electrodes placed on the thenar and hypothenar of the left hand, using a 50-Hz notch filter. An electrode attached to the left forearm served as ground for both ECG and skin conductance, which were recorded with the same amplifier (BrainAmp ExG, BrainProducts *GmbH*). In the “direct interaction” and “mediated interaction” conditions, data from observer and target were recorded synchronously by connecting all amplifiers to the same USB adapter feeding the data into BrainVisionRecorder (version 1.21.0102, BrainProducts GmbH).

### Quantification and statistical analysis Physiological data processing

#### EEG data

All pre-processing was done in EEGLAB, version 2020.0 (Delorme & Makeig, 2004), implemented in MATLAB R2019b (The Mathworks). Data were re-referenced to the right earlobe, and bipolar horizontal and vertical EOG channels were computed. Consistently bad channels (mean number = 2.01, range = 0 to 8 channels per participant and condition) and data segments with large artefacts were removed from the data (resulting in on average 1.6% of removed trials per participant and condition in the final epoched data). A bandpass filter was applied (finite impulse response filter, lower limit: 1 Hz, upper limit: 40 Hz, filterorder: 16500). Next, independent component analysis (ICA; implemented with the *runica* function in EEGLAB) was used for ocular artefact correction. Independent components that were clearly related to eye blinks or horizontal eye movements based on topography and time course were visually detected and removed (ranging from 2 to 6 components per participant). Afterwards, the weights of the remaining components were projected onto the original unfiltered data (Stropahl et al., 2018). Channels that had been removed before the ICA were interpolated (spherical interpolation); For some participants additional bad channels (mean number = 0.44, range = 0 to 3 channels per participant and condition) had to be interpolated. Data were then filtered with a bandpass filter with a lower limit of 0.2 Hz and an upper limit of 40 Hz (Finite impulse response filter, filter order = 16500, hamming window). Afterwards, data were segmented into stimulus-locked epochs of 4500ms lengths (1000 ms before and 3500 ms after stimulus onset) and baseline-corrected to 1000 ms before stimulus onset. A voltage threshold (between −70/70 μV and −100/100 μV) was manually set for each participant in a way that all trials with non-ocular artefacts were removed. The number of rejected trials varied from 0 to 31% per participant and condition and did not differ between conditions (M = 13% in all conditions).

For the time-frequency analysis, single-trial data of all electrodes were convolved with a complex Morlet wavelet as implemented in MATLAB (function *cwt* with parameter specification ‘cmor1-1.5’):

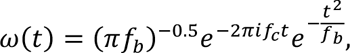

where *f_b_* = 1 is the bandwidth parameter, and *f_c_* = 1.5 is the wavelet center frequency. Specifically, for each participant, changes in time-varying energy were computed (square of the convolution between wavelet and signal) in the frequencies (1–40 Hz, linear increase) for the 1500 ms after shock onset. Power values were converted to decibels with respect to an average baseline from 500 to 50 ms before stimulus onset (Cohen, 2014). For analyses of peak-frequency power (see next paragraph), we subtracted the averaged data from 500 to 50 ms before stimulus onset as a baseline correction.

To analyse mu suppression, we determined the individual peak frequency of mu power for each participant by using the *restingIAF* toolbox (Corcoran et al. 2018). Although mu power has often been analysed as the mean of the power in the frequency range from 8-13 Hz, resting state oscillations over the somatosensory cortex often show a distinct peak in one specific frequency, which is suppressed upon somatosensory activation. The peak frequency differs between individuals (Berchicci et al. 2011; Gundlach et al. 2020). It is therefore useful to analyse desynchronization in individual peak frequencies to target oscillations functionally related to somatosensory activation, instead of averaging over a whole frequency band. Frequency peaks in the range from 8 to 13 Hz were detected in the baseline data from 1000 to 0 ms before shock onset in the “own pain” condition at central electrodes (C1, C2, C3, C4, C5, C6, CP1, CP2, CP3, CP4, CP5, CP6, FC1, FC2, FC3, FC4, FC5, FC6). For each detected frequency peak, the difference in average power between baseline (−1000 to 0 ms) and stimulation (0 to 1000 ms after shock onset) was calculated. The electrode and corresponding peak frequency with the strongest shock-related desynchronization for each participant was chosen for all further analyses. In 16 participants, a clear peak frequency was detectable. Peaks occurred at all possible frequencies except 9 Hz, and at the following electrodes: C1, C4, C3, CP2, CP3, C6, CP6, FC6, and FC2. The remaining 13 participants did not show a peak frequency, and for them data from the most frequent peak frequency and electrode were used (11 Hz at C4).

#### ECG data

ECG data were loaded into EEGLAB, filtered with a bandpass filter (lower: 1 Hz, upper: 30 Hz, finite impulse response filter, filterorder = 8250) and segmented into epochs of 2 s before and 8 s after stimulus onset for the stimulus-locked analyses. The MATLAB function *findpeaks* was used to detect the r-peaks in the segmented as well as the continuous data (for additional analyses of physiological coupling). Afterwards, data were visually screened for wrongly assigned or missing r-peaks. Data sections containing extrasystoles or otherwise undetectable R-peaks were treated as missing values. The interbeat interval (IBI) in ms was calculated for every pair of heartbeats and used as IBI value for each original data point in between the two heartbeats. In this way, the IBI trace had the same time resolution as the original data. For the analysis of condition differences in IBI responses to shocks, the mean IBI from 2 to 5 s after cue onset (Sperl et al. 2016) was computed and baseline-corrected to the mean of the 2 s before cue onset.

#### Skin conductance data

Skin conductance responses (SCRs) were analysed using the Ledalab-Toolbox (Benedek & Kaernbach, 2010, version 3.4.8) in Matlab. Data were downsampled to 50 Hz, smoothed with an adaptive gaussian window and visually screened for strong artefacts, which were spline-interpolated. Afterwards, a continuous deconvolution analysis was conducted to separate phasic from tonic skin conductance activity (Benedek & Kaernbach, 2010). In the following, the mean phasic driver activity 1–4 s after shock onset was used for analyses (for targets’ SCR in physiological coupling). For the observer data (condition differences and physiological coupling), four different time lags (1–4 s) were considered to account for a time lag in the observer’s response to the target’s pain expression.

### Statistical analyses

We first outline the general statistical analysis approach before specifying the details. One set of statistical analyses examined the observers’ responses to the targets’ pain (empathic accuracy, unpleasantness ratings, neural and physiological responses; see Fig. 1B, left side). In these analyses, the predictors, shock intensity, condition (direct vs. mediated) and their interaction were tested. A significant effect of shock intensity indicates that the observer’s responses are influenced by the other’s pain and are therefore interpreted as empathic. A significant effect of condition indicates that social presence generally changes the observers’ behaviour and physiology, whereas an interaction between shock intensity and condition indicates that social presence alters the sensitivity to another’s pain. The second set of analyses examined physiological coupling between observer and target responses (see Fig. 1B, bottom). In these analyses, the targets’ responses, the condition (direct vs. mediated) and their interaction served as predictors. A significant effect of targets’ responses indicates that observers’ and targets’ responses are generally coupled, whereas an interaction with condition indicates stronger coupling in one condition compared to the other. In both sets of analyses, (generalized) linear mixed models with single trials (Level 1) nested in observers (Level 2) and observers nested in targets (Level 3) were calculated. For theta activity, responses were averaged over trials. In cases where the hypothesized effects were not significant, Bayes Factors were calculated by comparing the Bayesian Information Criterion between models with and without the predictor in question. This was done to confirm evidence for the null-hypothesis (Lee and Wagenmakers 2014). For the peak-mu analysis, permutation tests over the whole time course were conducted, as there was no predefined time window. To test for effects of intensity, condition and their interaction beyond mu suppression, exploratory permutation tests on the whole EEG data space were conducted.

To assess the robustness of the findings, the analyses of mu, IBI and SCRs were repeated with data averaged over trials. In these analyses, the factor intensity was dichotomized into low and high intensity (low: 1 to 10; high: 11 to 20). The results of these analyses are not reported, as they did not differ from the single-trial results. In the figures, data are dichotomized into low and high intensities for display purposes only. Permutation tests were carried out in MATLAB (version R2019b, The Mathworks), and all other statistical analyses were carried out in R (version 4.0.2, *R core team*).

#### Behavioural data

To test for condition differences in empathic accuracy, we conducted a negative binomial generalized linear mixed model (using the function *glmer.nb* in the *lme4* package in R, Bates et al., 2020) on observers’ single-trial ratings of the other’s pain (Lawless 1987). To find the best random slopes structure, first models with the full fixed-effects structure and different random slopes were compared using the Akaike Information Criterion (AIC, Akaike, 1998). An AIC difference greater than 2 was set as the threshold for a significant difference (Burnham and Anderson 2004). Then one fixed predictor after the other was added to the model with the optimized random-effects structure, and only predictors that significantly improved the model were kept. The same procedure was used to test for differences in unpleasantness ratings between the conditions.

#### EEG data

To test whether mu suppression was modulated by shock intensity (20 levels), condition (direct vs. mediated) or their interaction, permutation tests were conducted on the mu peak-frequency power within the window from −500 to 1500 ms after stimulus onset (see Cohen, 2014). To test for effects of shock intensity on mu suppression, Spearman correlations between normalized single-trial power and single-trial shock intensity were calculated for each data point and across participants. This procedure yielded a time course of correlation coefficients between peak-frequency power and shock intensity. Correlation values were z-transformed by comparing them to a permutation-based null-hypothesis distribution (based on randomly shuffling over trials 1000 times). To correct for multiple comparisons, a maximum value correction was used (Cohen, 2014).

To test for condition differences in mu suppression regardless of shock intensity, mu power was averaged over trials and compared between conditions. For the null-hypothesis distribution, the assignment of condition was randomly shuffled over participants (1000 permutations). To test for condition differences in the correlation between shock intensity and power (whether power tracked shock intensity to a larger degree in the direct condition), the condition difference between correlation coefficients was calculated for each data point and participant and compared to a random distribution (shuffled between conditions in 1000 permutations). For a comparison, we also analysed power in the traditional mu band (8–13 Hz, electrode C4, 500–1000 ms after shock onset). We chose the time window according to previous literature and to be comparable to the time window in which the response to own pain occurred (Zebarjadi et al. 2021).

For the effects of condition, shock intensity and their interaction on theta responses, we analysed power in the traditional theta band (4–8 Hz, electrode Fz, 0–500 ms after shock onset). We chose the time window according to previous literature (Mu et al. 2008; Peng et al. 2021). In the latter two analyses, we used a linear mixed model with condition averages nested in observers and observers nested in targets and dichotomized shock intensities.

Finally, in an exploratory analysis, using the same permutation test method as described above for peak-frequency mu, we looked for main effects of shock intensity (20 levels), condition and their interaction on power across the whole time-frequency-electrode space (1–20 Hz, all electrodes). For the effect of shock intensity, a time window from 0 to 1500 ms was used, for the other analyses a time window from −500 to 1500 ms. For the effect of shock intensity data were downsampled to 125 Hz to reduce computation time. Again, a maximum value correction and additionally a cluster size correction (Cohen, 2014) were used to correct for multiple comparisons.

#### IBI and SCRs

To test for condition and shock intensity (20 levels) effects on the observers’ IBI, linear mixed models on the singletrial IBI averages were conducted (using the *lmer* function in *lme4* in R). The same procedure was used for the skin conductance data. Here, first the time lag with the greatest SCR over all conditions was selected for further analyses. To account for interindividual variation in SCR levels, SCR means were normalized by dividing them by participants’ individual standard deviation. As SCR data were not normally distributed, generalized linear mixed models (using the *glmer* function in *lme4*, Gamma family, log-link function) were used for these data. To assess the robustness of the findings, all analyses were also carried out with data averaged over trials (with dichotomized shock intensities).

To test for coupling between targets’ and observers’ IBI and SCRs, similar (generalized) linear mixed models were used, but this time the targets’ IBI response or SCR was entered as a fixed predictor instead of the stimulus intensity. For all (generalized) linear mixed models, fitting of random- and fixed-effects structure was carried out in the same way as for the single-trial behavioural data. To assess the robustness of the findings, skin conductance coupling was also analysed by calculating Spearman’s correlation coefficients between targets’ and observers’ SCRs and comparing them between conditions using a linear mixed model. Correlations were calculated for the four time lags, and the time lag with the highest correlation coefficients across both conditions was chosen for the linear mixed model. For the robustness analysis of the IBI coupling, the Spearman’s correlation between the targets’ and the observers’ continuous IBI over the whole task was calculated. In this analysis, IBI data were smoothed with a moving average function of 2 seconds to reduce the influence of strong outliers. The correlation coefficients were calculated for 20 different time lags (steps of 0.5 s) between target and observer data. For testing condition effects, the lag with the highest correlation over all conditions was used. Correlation coefficients were transformed using the Fisher’s z-transformation to obtain normally distributed data. They were then compared between conditions using linear mixed models with correlation coefficients from different conditions (Level 1) nested in observers (Level 2) and targets (Level 3). The results of these analyses are reported in the results section when they diverge from the results of the singletrial analyses.

### Control analysis of target expressivity

Empathic accuracy depends on the expressivity of the other (Zaki et al. 2008), and differences between direct and mediated interactions might result from altered expressivity of the targets in either condition. To test whether the targets show systematic differences in their pain expression between the conditions, a control experiment with a different sample was conducted. Thirty-one participants (25 females, 6 males, mean age(SD) = 23.51(4.70)) were shown 100 segments from the video recordings of both direct and mediated interaction without being aware of that manipulation. Video segments included four seconds before and four seconds after an electric shock and were chosen such that there was an equal number of videos from each original session, condition and target pain rating (summarized in 5 bins of 20 rating points each). Videos were shown in random order. After each video the participants had 10 seconds to rate how painful the stimulus was for the target in the video on a visual analogue scale ranging from “not painful at all” to “extremely painful”. If targets expressed their pain differently in the two conditions, we would expect condition differences in the mean pain ratings or in the empathic accuracy of the control participants. Mean pain ratings and mean empathic accuracy scores (Spearman’s correlations) were then compared between the conditions using t-tests.

### Pre-registration

A pilot study using a similar design was pre-registered at OSF: https://osf.io/gcyqs. Behavioral and physiological raw data as well as the main analysis code have been deposited at https://osf.io/pqmra/?view_only=df6b8e7cd19743d481d86ef7cb83cb83 and are publicly available at the date of publication. Further data and code are available upon reasonable request from the first author.

## Results

### Social presence does not influence empathic accuracy

To assess effects of social presence on empathic accuracy, observers’ ratings of targets’ pain were predicted from shock intensity, condition (“direct” vs. “mediated”) and their interaction using generalized linear mixed models with single trials nested in observers and observers nested in targets. The best model had a random-effects structure containing random slopes for *condition* and *intensity* (AIC difference to the next best model during fitting of random effects: −63). Moreover, it contained a fixed effect of *intensity* (b(SE) = 0.07(0.01), *z* = 10.24, *p* < 0.0001, AIC difference to a model without the fixed effect of intensity: −14). The Bayes factor for the hypothesized interaction of condition and intensity was 0.075, indicating strong evidence for the model without the interaction (Lee and Wagenmakers 2014). These results show that the intensity of the shocks received by the targets predicted the observers’ pain ratings (Fig. 2A & B), indicating meaningful empathic accuracy. However, this effect did not differ between conditions. To control for possible learning effects, we assured that participants were not more accurate in the second condition (interaction between time and intensity on observer ratings: b(SE) = 0.001(0.005), *z* = 0.21, *p* = 0.84), and that the order of conditions did not moderate any effects of social presence on empathic accuracy (interaction between intensity, condition, and order condition: b(SE) = −0.002(0.024), *z* = −0.073, *p* = 0.942).

**Figure 2.**
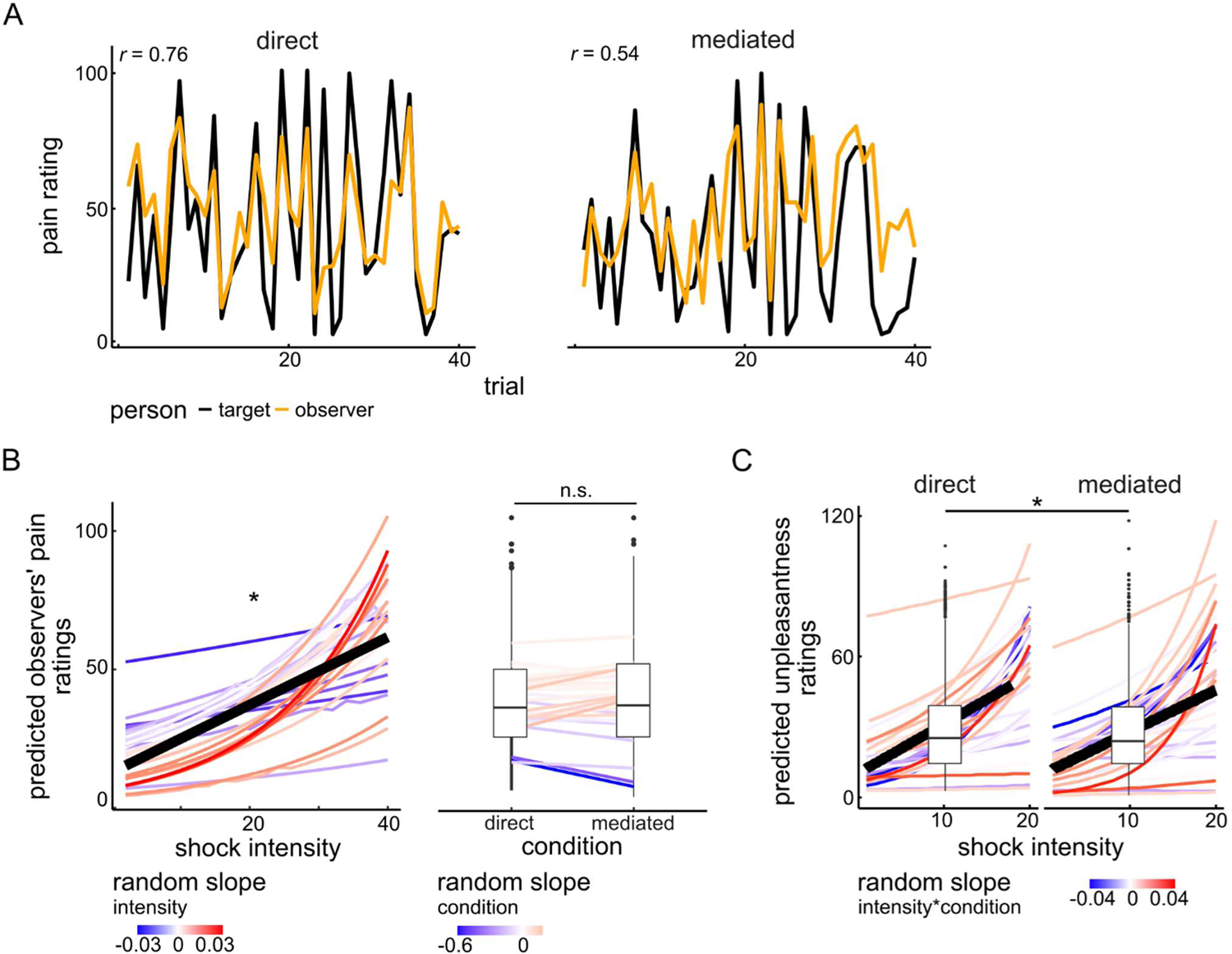
(A) Example pain rating data over trials from one randomly chosen sample dyad. (B) Predicted observer pain ratings averaged over direct- and mediated-interaction conditions. The black thick line represents the fixed effect of shock intensity; single coloured lines represent predicted data from single participants. The colour shading from blue to red represents the value of the random slope for intensity of each participant (left). Predicted observer pain ratings depending on condition (direct vs. mediated). Lines represent values for single participants. The colour shading from blue to red represents the value of the random slope for condition of each participant. The boxplots represent summary statistics for each condition (C) Predicted observer unpleasantness ratings in direct and mediated interaction. The black thick line represents the fixed effect of shock intensity; single coloured lines represent predicted data from single participants. The colour shading from blue to red represents the value of the random slope for the interaction between intensity and condition of each participant. The boxplots represent summary statistics for each condition. * = *p* < 0.01, n.s. = not significant.

As the familiarity within the dyads varied slightly, we did additional control analyses to assure that the varying familiarity between targets and observers in our sample did not masks effects of social presence on empathic accuracy (Ma et al. 2011; Meyer et al. 2013; Motomura et al. 2015). Although familiarity might modulate the effect of social presence on empathy (see supplementary materials for detailed analyses), the results show that it is unlikely that the inclusion of familiar dyads masked any effects of social presence on empathic accuracy, rather the contrary. However, it has to be noted that the number of dyads who had met more than twice was very small (four dyads) and therefore results of these analyses have to be interpreted with caution.

### Social presence increases unpleasantness ratings

We used a generalized linear mixed model to assess whether shock intensity and condition (“direct” vs. “mediated”) predicted the unpleasantness ratings. The best model had a random-effects structure containing random slopes for the interaction between *condition* and *intensity* (AIC difference to the next best model during fitting of random effects: −29). It contained a fixed effect of *intensity* (b(SE) = 0.07(0.01), *z* = 8.07, *p* < 0.0001) and a fixed effect of *condition* (b(SE) = −0.14(0.05), *z* = −2.63, *p* = 0.009, AIC difference to a model without the fixed effect of *condition*: −4). The Bayes factor for the hypothesized interaction of condition and intensity was 0.021, indicating very strong evidence for the model without the interaction (Lee and Wagenmakers 2014). This shows that the intensity of the shocks received by the targets predicted the observers’ unpleasantness ratings (Fig. 2C), but equally so in both conditions. However, the observers rated the pain of the target as slightly more unpleasant in the direct than in the mediated interaction (mean difference (SD) = 1.47(6.45)).

### Mu suppression is not sensitive to observed pain

Using a permutation test, we analysed whether mu suppression was sensitive to the observed pain intensity, and whether it was reduced in the mediated condition. For each participant, we chose an individualized frequency (in the range of 8 – 13 Hz) and electrode (of central electrodes), which showed the strongest desynchronization in the own pain condition. Data from 29 dyads were included in these analyses, as data from the own pain condition of one participant were not analysable due to technical issues during data collection. The analyses of peak mu suppression in the “direct” and “mediated interaction” conditions showed no effect of *intensity* or *condition,* nor an interaction that was significant after maximum value correction (see Fig. 3A & 3B middle and right). These results show that there was no mu suppression which was sensitive to others’ pain intensity in either condition. Similarly, analysing averaged power over the canonical mu band (8–13 Hz) yielded no significant effect of either factor (*intensity*: b(SE) = 0.34(0.30), *t*(df) = 1.13(87), *p* = 0.26; *condition*: b(SE) = −0.04(0.30), t(df) = −0.14(87), *p* = 0.88; *interaction intensity*condition*: b(SE) = −0.02(0.42), *t*(df) = −0.04(87), *p* = 0.97). However, peak mu significantly differed between pain levels when participants experienced pain themselves (Fig. 3A left).

**Figure 3.**
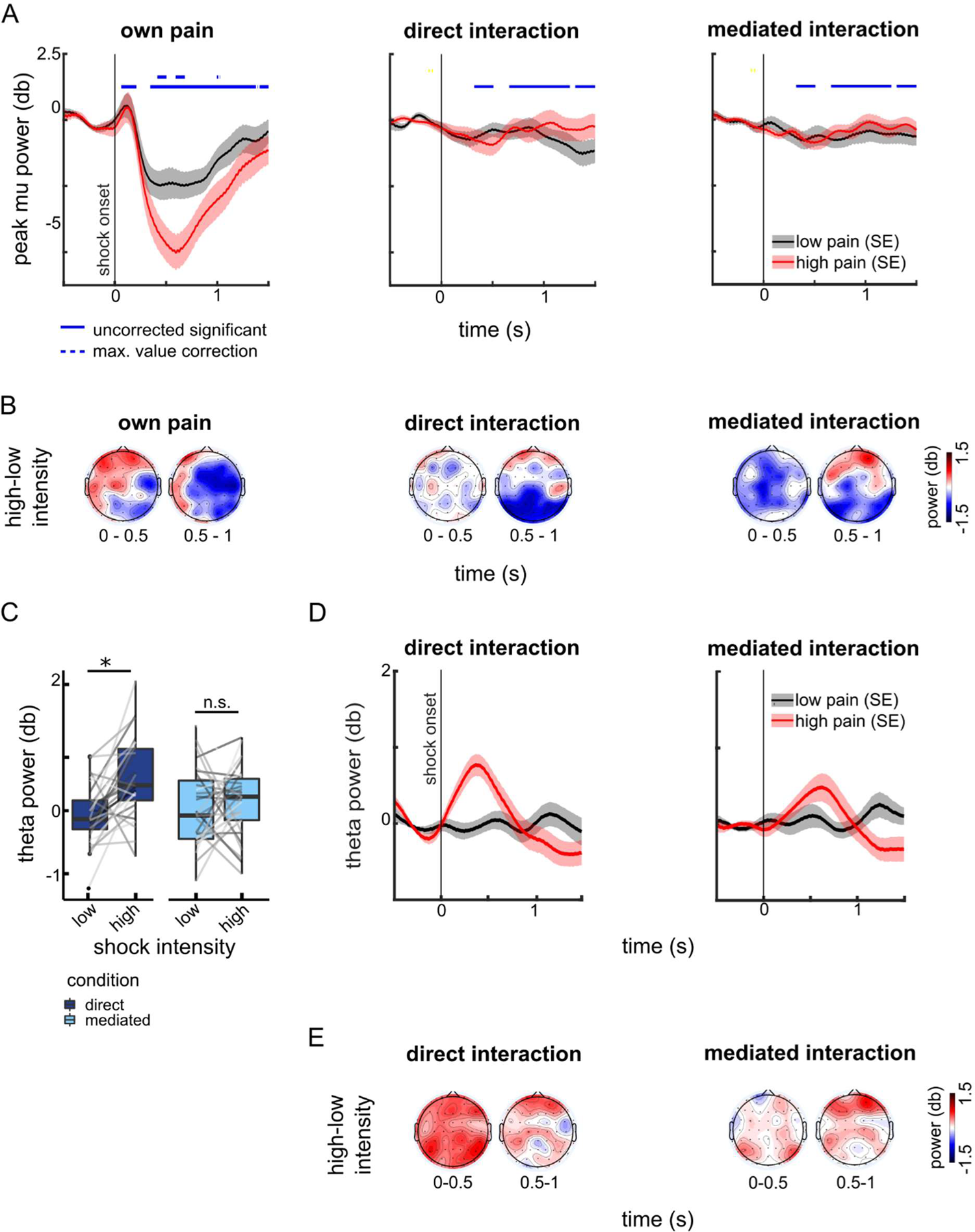
(A) Shown is peak mu power averaged over participants, dichotomized into low- and high-intensity shocks for display purposes only. Blue solid lines indicate time windows where the main effect of intensity reached significance (uncorrected level); blue dotted lines indicate time windows where the intensity effect survived maximum value correction. The main effect of condition and the interaction of condition x intensity did not survive maximum value correction, at any time-point. On the left side, the clear effect of pain intensity on mu power in the own-pain condition can be assessed. (B) Shown is the topography of averaged peak mu power differences between high- and low-intensity shocks in the three conditions. (C) Shown are boxplots for theta power averaged over 4–8 Hz and 0–500 ms after shock onset at electrode Fz for the direct and mediated conditions. Grey lines depict means from single participants. Significance asterisks refer to post-hoc tests from linear mixed models. * = *p <* 0.001.(D) Shown is the time course of averaged theta power (4–8 Hz) after stimulus onset. (E) Shown is the topography of averaged theta power differences between high- and low-intensity shocks in the two conditions.

### Sensitivity of theta power to others’ pain is modulated by social presence

We then tested whether theta responses were sensitive to observed pain intensity, and whether their sensitivity was modulated by social presence. Data from 30 dyads were included in these analyses. The best-fitting linear mixed model for the power averaged over the canonical theta band (4–8 Hz) yielded a significant main effect of *intensity* (b(SE) = 0.57(0.14), *t*(df) = 3.99(90), *p* < 0.001) and a significant interaction between *condition* and *intensity* (b(SE) = −0.46(0.20), *t*(df) = −2.30(90), *p* = 0.024, AIC difference to next best model: 2.9). Follow-up models on the significant interaction showed a significant effect of *intensity* in the direct condition (b(SE) = 0.57(0.13), *t*(df) = 4.46(28.99), *p* < 0.001), but not in the mediated condition (b(SE) = 0.10(0.15), *t*(df) = 0.70(28.99), *p* = 0.49). This indicated that frontal theta was more responsive to the other’s shock intensity in the direct condition than in the mediated condition (see Fig. 3C, D, E).

### Exploratory analyses of whole time-frequency-electrode space

Grand Averages of EEG power in the different conditions can be found in supplementary figure S1-S6. To test for modulations of neural activity by observed shock intensity or social presence beyond mu and theta band, we ran a cluster based permutation test on the whole time-frequency-electrode data space. We used a cluster size correction and a maximum value correction to correct for multiple comparisons. Data from 30 dyads were included in these analyses. During both direct and mediated interaction, others’ pain *intensity* was positively related to 2 to 6 Hz power between 56 and 840 ms after shock onset at the frontal and central electrodes (cluster 1, maximal at F4, see Fig. 4A left, B and E top left). Others’ pain *intensity* was also negatively related to 12 to 20 Hz power between 680 and 1040 ms after shock onset over parietal and occipital electrodes (cluster 2, maximal at P2, see Fig. 4A left, B and E bottom left).

**Figure 4:**
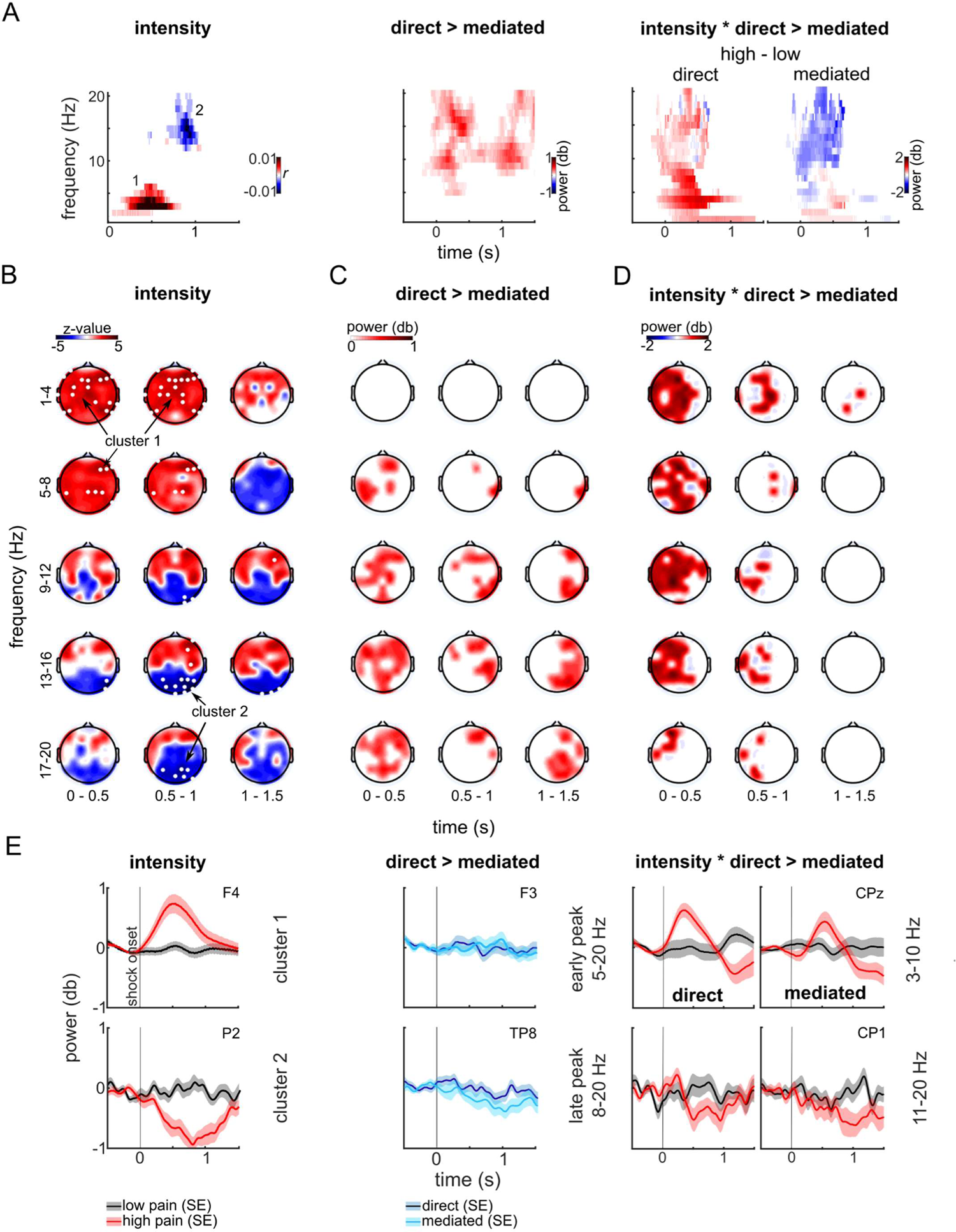
(A) Clusters found in the permutation tests on the correlation between shock intensity and EEG power across both conditions (left), on the main effect of condition (middle) and on the interaction between shock intensity (dichotomized for display purposes only) and condition (right), averaged over electrodes. (B) Results of the permutation test on the correlation between shock intensity and EEG power across direct and mediated conditions. Data points that were significant after cluster size correction are displayed in colour; data points that were significant after maximum value correction are marked in white. (C) Results of the permutation tests on the condition effect. Data points of the largest cluster (uncorrected significant) are displayed in colour. (D) Results of the permutation tests on the interaction between condition and intensity (dichotomized for display purposes only). Data points of the largest cluster (uncorrected significant) are displayed in colour. (E) Power time course of cluster 1 (top left) and cluster 2 (bottom left), and power time course for the early cluster (5–20 Hz, F3, middle top) and the late cluster (8–20 Hz, TP8, middle bottom) showing a condition effect and power time course of lower frequencies (3–9 Hz; top, right), and higher frequencies (10–20 Hz; bottom, right) for interaction effects between condition and intensity. Data are dichotomized into low and high intensities for display purposes only.

Neither a permutation test on the *condition* effect (mediated vs. direct) nor the interaction between *condition* and *intensity* yielded any cluster that survived the cluster size correction or the maximum value correction. However, due to the exploratory nature of the analyses, we further examined the biggest cluster, which was significant on an uncorrected level. In direct compared to mediated interactions (main effect of condition; see Fig. 4A middle, C and E middle), theta/alpha (5–12 Hz) power at the frontal electrodes and lower beta (13–20 Hz) power at the frontal, central and centro-parietal electrodes was enhanced in an early time window (−256–644 ms after shock onset). In a later time window (511–1500 ms after shock onset), alpha/lower beta (8–20 Hz) power at the centro-parietal electrodes was enhanced during direct interactions. For the interaction between *condition* and *intensity*, the biggest uncorrected significant cluster showed a stronger positive effect of *intensity* in the “direct” than in the “mediated” condition. This cluster spanned 3–20 Hz between −220 and 1380 ms after shock onset and included most left-hemisphere and central electrodes (see Fig. 4A right, D). Its time course is displayed separately for low (3–9 Hz, electrode with maximal interaction: Cz, Fig. 4E top right) and high frequencies (10–20 Hz, electrode with maximal interaction: CP1, Fig. 4E bottom right).

### Social presence does not modulate physiological responses to other’s pain

Next, we tested whether observers showed stronger cardiac responses (interbeat interval, IBI, responses) to observed pain in the “direct” than in the “mediated” condition, again using linear mixed models on single trials. The best model contained random slopes for *condition* and *intensity* but no fixed effects. The Bayes factor for the hypothesized effect of intensity was 0.027, indicating very strong evidence for the model without the effect. The Bayes factor for the hypothesized interaction of condition and intensity was 0.088, indicating strong evidence for the model without the interaction (Lee and Wagenmakers 2014). These results indicate that observers’ IBI responses were not sensitive to the observed shock intensity on the sample level (see Fig. 5A).

**Figure 5:**
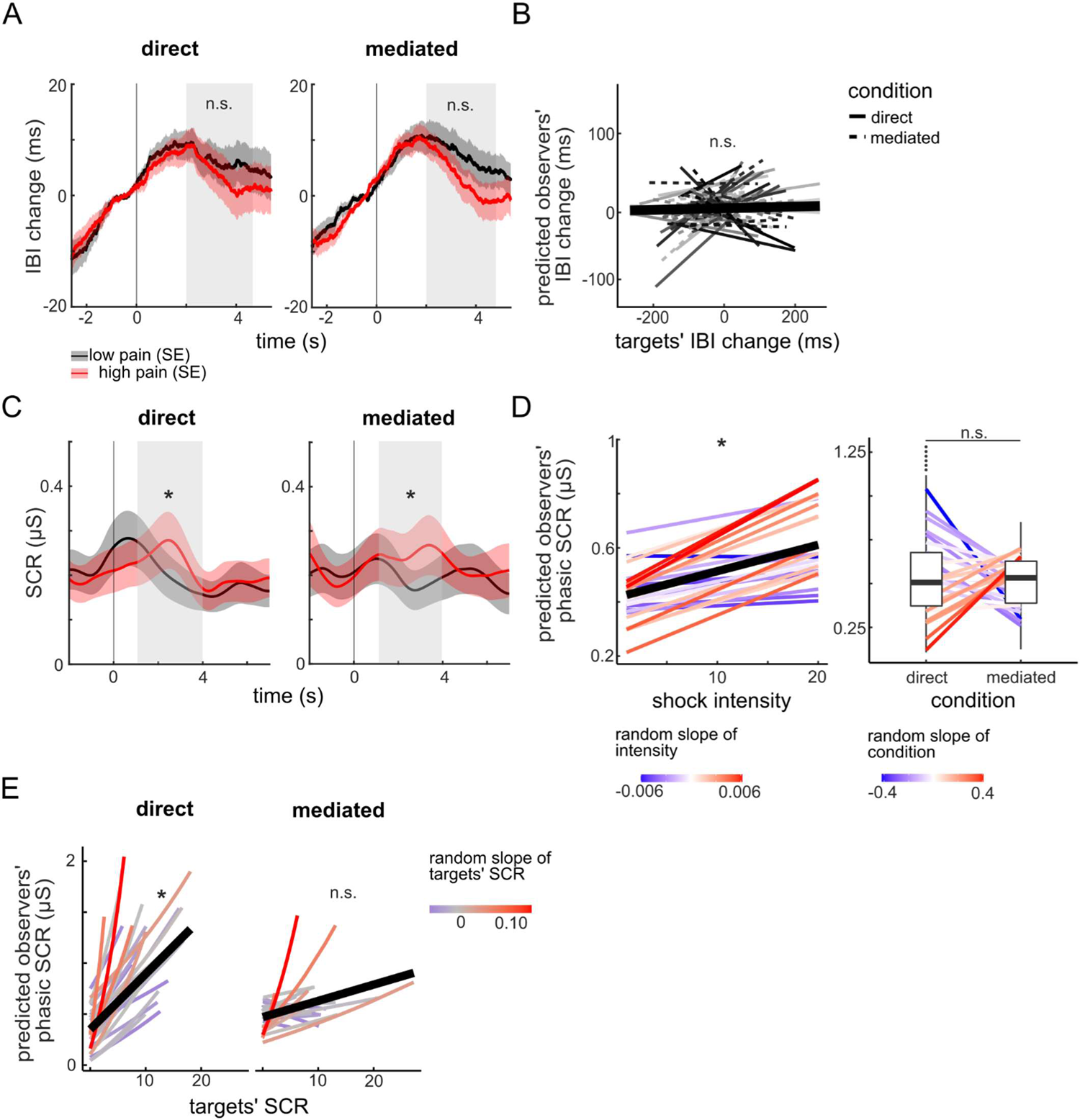
(A) Grand averages of observers’ IBI responses in direct and mediated interactions, dichotomized into low- and high-intensity trials for display purposes only. (B) Observer IBI responses predicted from the single-trial model on physiological coupling. The thick black line represents the fixed effect of target IBI; the thin lines represent predicted data for single participants. (C) Grand averages of observers’ SCRs in direct and mediated interactions, dichotomized into low- and high-intensity trials for display purposes only. (D) Observer SCRs predicted from shock intensity in the generalized linear mixed model. The black thick line represents the fixed effect of intensity; the coloured lines display predicted values for single participants. The colour shading from blue to red represents the value of the random slopes of shock intensity for each participant (left). Observer SCrs predicted from condition. The coloured lines represent values for single participants, the colour shading from blue to red depicts the random slope of condition for the single participants (right). (E) Observer SCRs predicted from target SCRs in the generalized linear mixed model on physiological coupling. The black thick line represents the fixed effect of targets’ SCRs; the coloured lines display predicted values for single participants. The colour shading from blue to red represents the value of the random slopes of target SCR for each participant. * = *p* < .001, n.s. = not significant.

Moreover, we tested whether observers’ skin conductance responses (SCRs) were more responsive to observed pain intensity in the “direct” than in the “mediated” condition. Observers’ SCRs were greatest in the time window from 1 to 4 seconds after shock onset, hence this time window was used for all further analyses. The best single-trial linear mixed model on observers’ SCR to the observed shocks contained random slopes for *condition* and *intensity* and a fixed positive effect of *intensity*. The Bayes factor for the hypothesized interaction of condition and intensity was 0.036, indicating strong evidence for the model without the interaction (Lee and Wagenmakers 2014). These results indicate that observers’ SCRs were sensitive to the shock intensity, but equally so in direct and mediated conditions (Fig. 5C and 5D). All model parameters are listed in Table 1. Models with dichotomized intensities yielded the same results.

**Table 1:**
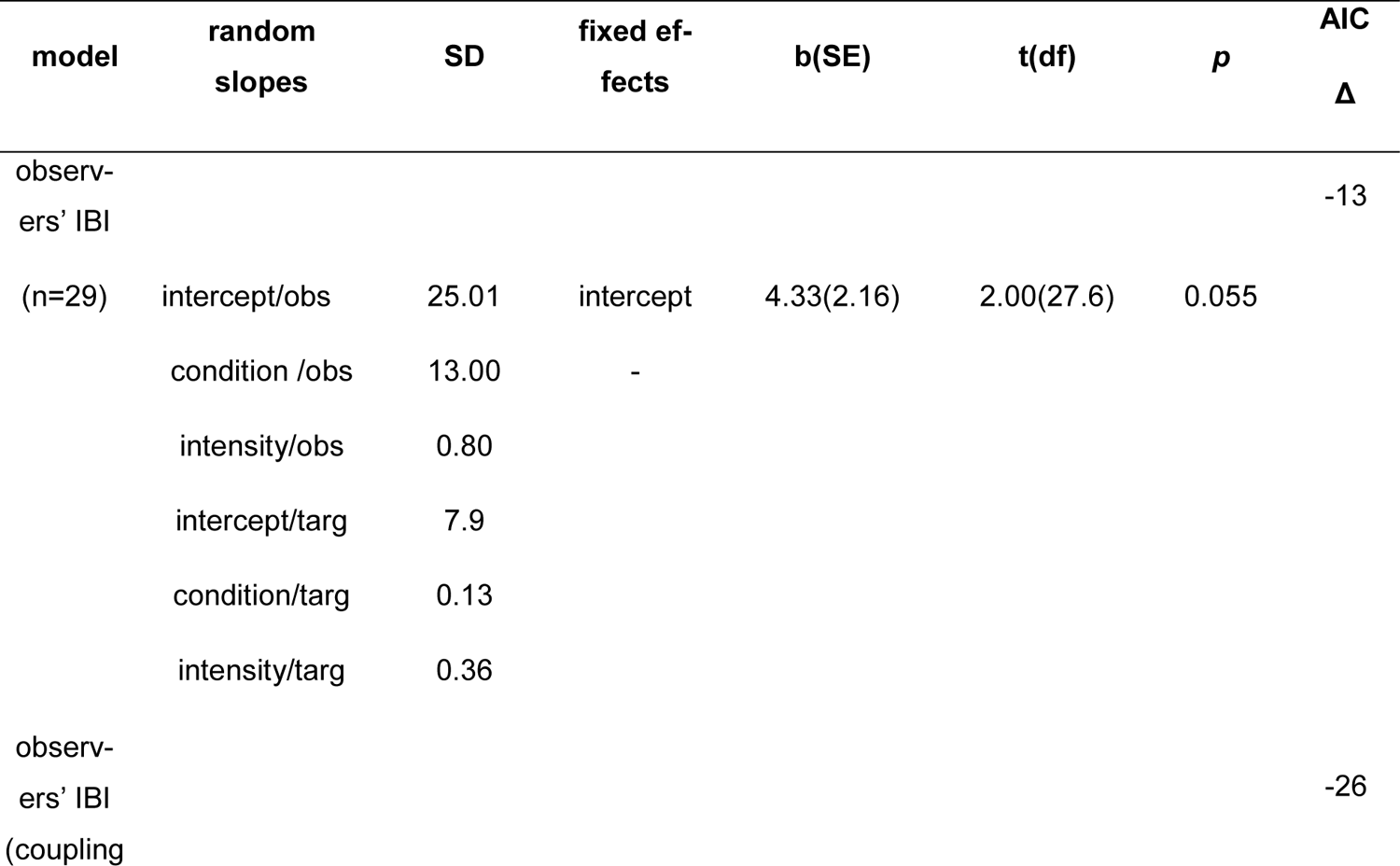
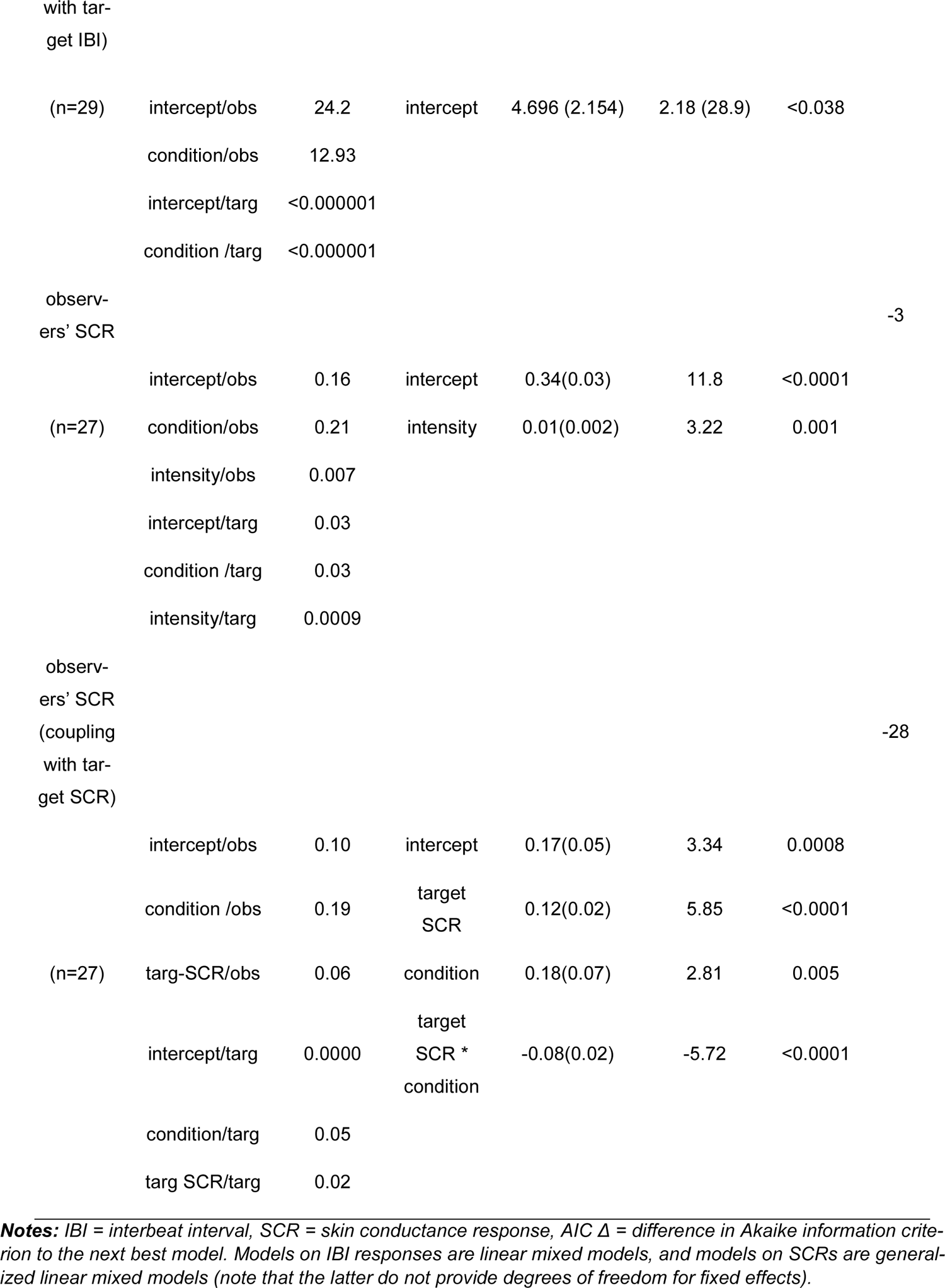
Results of the single-trial (generalized) linear mixed models on IBI responses and SCRs

### Social presence enhances skin conductance coupling between target and observer

Finally, we used a single-trial linear mixed model to test whether the direct interaction enhanced the coupling between targets and observers IBI responses. To this end, the model tested whether observers’ IBI responses could be predicted from targets’ IBI responses and whether this effect was modulated by the condition (“direct” vs. “mediated” interaction). The best model contained a random slope for *condition* but no fixed effects. The Bayes factor for the hypothesized effect of targets’ IBI was 0.059, indicating strong evidence for the model without the effect. The Bayes factor for the hypothesized interaction of condition and targets’ IBI was 0.076, indicating strong evidence for the model without the interaction (Lee and Wagenmakers 2014). These results indicate that there was no coupling between targets’ and observers’ IBI responses (Fig. 5B). Comparing the correlations between the continuous IBI traces in the two conditions showed the same results (correlations were highest for a lag of 2 seconds).

The same rationale was used to test for effects of condition on coupling between targets’ and observers’ skin conductance responses to pain. The best single-trial linear mixed model on SCR coupling – predicting observers’ SCR from targets’ SCR – contained a random slope for *condition* and *target SCR* and fixed effects of *target SCR*, *condition* and their interaction (for model parameters, see Table 1). Follow-up models on the interaction between *condition* and *target SCR* showed a significant positive effect of *target SCR* on observers’ SCR in the “direct” condition (b(SE) = 0.12(0.03), *t* = 4.15, *p* < 0.0001), but not in the “mediated” condition (b(SE) = 0.03(0.02), *t* = 1.64, *p* = 0.1). When comparing the correlation coefficients of targets’ and observers’ SCRs between conditions, there was only a marginally significant effect of *condition* (b(SE) = −0.07(0.04), *t*(df) = −1.81(26), *p* = 0.082). These results indicate that coupling between targets’ and observers’ SCRs was greater in the direct than the mediated interaction, but that the effect was rather small (see Fig. 5E).

### Targets do not express their pain differently in the two conditions

To test whether possible differences in empathic responses towards the targets’ pain in the two conditions were due to different expressivity of the targets’ in the two conditions, we conducted a control experiment. We showed 100 video segments of targets’ experiencing pain from the “direct” and “mediated” conditions to 31 naïve participants, who rated the pain intensity of the shocks displayed in the video. These participants were unaware of the condition manipulation in the videos. If targets’ expressed their pain e.g. systematically stronger in the “direct” than in the “mediated” condition, the new pain ratings should be higher for videos from the “direct” than from the “mediated” condition. This was not the case. The means of the pain ratings from the video control experiment did not differ significantly between conditions (mean difference = 0.79, *t*(30) = 1.66, *p* = 0.11, Cohen’s d = 0.3). The mean Spearman’s correlations between video observers’ ratings and shock intensity also did not differ between videos from direct and mediated interaction (mean_direct_(SD) = 0.36(0.16), mean_mediated_(SD) = 0.32(0.13), mean difference = 0.040, *t*(30) = 1.36, *p* = 0.18). These results indicate that the targets did not express their pain significantly differently in the direct versus mediated interaction.

## Discussion

Although mediated social interactions through video calls are becoming the new norm, the impacts on understanding others and their feelings have not yet been researched thoroughly (Grondin et al. 2019), especially in social neuroscience. In the current study, we explored how a video-mediated interaction affects empathy for pain on behavioural, physiological and neural levels. We expected that less availability of social cues in a mediated interaction would hamper empathizing with the other. However, we found that observers were just as accurate in judging the other’s pain in the mediated as in the direct interaction. Moreover, participants experienced the other’s suffering as only slightly less unpleasant in the mediated interaction. On the neural level, we did not find mu suppression over the somatosensory cortex that was sensitive to the other’s pain in either condition. However, mid-frontal theta tracked the other’s pain intensity more in the direct than in the mediated interaction. Exploratory analyses of the whole time-frequency-electrode space showed no additional differences between direct and mediated conditions after correcting for multiple comparisons. On a physiological level, observers’ SCRs were coupled to targets’ SCRs to a stronger degree in the direct compared to the mediated interaction. In sum, behavioural empathy was not reduced in the mediated interaction, whereas some neural and physiological aspects of empathy were dampened.

### Effects of social presence on behavioural aspects of empathy for pain

Surprisingly, among the many studies on empathy for pain, hardly any measured empathic accuracy, and none explored which type of information is necessary or sufficient to judge others’ pain accurately (Gauthier et al. 2008; Leonard et al. 2013; Laursen et al. 2014). In story-based empathy studies, empathic accuracy for emotion was reduced when auditory linguistic information was completely removed, whereas missing visual information did not impact empathic accuracy (Zaki et al. 2009; Jospe et al. 2020). Similarly, we reveal that participants could judge the targets’ pain quite well, and this ability did not decline in the mediated interaction. In contrast to the story-based paradigm, our results imply that visual information (apparent in both the direct interaction and the video calls) is sufficient for empathic accuracy for others’ pain. As our participants did not experience severe pain (expressed by moaning or crying), auditory information might have been less important than for example in empathic responses to the pain of hospital patients (Agahi and Wanic 2020). As our control analysis showed no condition differences in target expressivity, we can be assured that these did not mask true condition differences in empathic accuracy.

Although observers showed more affective empathy in the direct than in the mediated condition, the effect size was so small that it was practically negligible. One reason for that might be that many of our participants remembered their own recent pain experience to help them feel with the similar experience of the other, as they reported in the debriefing questionnaires. This strategy might have led to imagination of others’ pain independent of the medium, causing similar affective empathy (Goubert et al. 2005).

### Effects of social presence on neural aspects of empathy for pain

As most former studies on empathy for pain used abstract cues or static pictures, we expected to find even stronger mu suppression in our paradigm using real stimuli and focusing on individual peak-mu frequency (i.e., Perry, Bentin, et al. 2010; Riečanský and Lamm 2019; Zebarjadi et al. 2021). Instead, we did not find any mu suppression in response to others’ pain. Speculatively, mu suppression is a compensatory mechanism that aids empathy for pain via somatosensory representation of others’ pain if insufficient sensory cues are available. Alternatively, it might be specific for empathy for pain that is caused by visually perceivable physical injury, and might not be sensitive to facial expressions of pain. Otherwise, it may be a weak signal that requires many repetitions – and stronger pain signals – to find a significant effect. However, if mu suppression is a general empathy mechanism, our analysis should have been sensitive enough to detect it. Therefore, our null findings on mu suppression align with recent criticism of its robustness and validity as a mechanism underlying empathy in general (Hobson and Bishop 2016).

The mid-frontal theta/delta response constitutes another neural component of empathy for pain that has so far been rarely examined in EEG studies (but see Mu et al. 2008; Peng et al. 2021). Mid-frontal theta has been related to the salience, unexpectedness and aversiveness of many different types of stimuli (González-Roldan et al. 2011; Güntekin and Başar 2014; Cavanagh and Shackman 2015). Therefore, the heightened sensitivity of theta to the other’s pain level might indicate that the other’s pain elicits more arousal and negative affect in the direct interaction (Balconi et al. 2009), although without resulting in measurable behavioural differences. Mid-frontal theta might stem from the ACC, which shows reliable activity to both own and others’ pain in fMRI studies (Cavanagh and Frank 2014; Fallon et al. 2020). Confirming this assumption with source analyses was beyond the scope of the current paper but could be an important step in future studies. The exploratory single-trial permutation test confirmed on a trend level the interaction between others’ pain intensity and direct versus mediated interaction.

Lastly, our exploratory analysis revealed stronger parietal beta suppression relating to stronger observed pain irrespective of the condition. Parietal beta decrease has been linked to attention to affective touch (von Mohr et al. 2018). Future EEG studies should clarify its role in empathy for pain.

### Effects of social presence on physiological coupling

Observers’ cardiac activity was not sensitive to others’ pain and showed no coupling with targets’ cardiac activity. In contrast, previous empathy studies found cardiac coupling in emotional empathy (Zerwas et al., 2021), especially when semantic and auditory information was missing (Jospe et al., 2020). The shocks elicited a strong cardiac acceleration in the targets in our paradigm (data not shown in this paper). Possibly these responses were not easily mimicked by observers cardiac activity, which might be one reason for the discrepancies with former studies (Goldstein et al. 2017).

In contrast, we found coupling in skin conductance responses, which was the one aspect of pain empathy that was markedly reduced in the mediated interaction. This finding indicates that the physiological coupling component of empathy, specifically in SCR, might rely on physical proximity (Chatel-Goldman et al. 2014; Murata et al. 2020) and possibly olfactory cues that are missing in mediated interactions (de Groot et al. 2014; Calvi et al. 2020).

### Conclusions and future directions

Many recent studies examined direct interactions between participants, claiming that it is necessary for understanding social cognition (Redcay and Schilbach 2019; Fan et al. 2021; Levy et al. 2021). However, few studies have explicitly compared these new paradigms to similar tasks using mediated interactions (but see e.g. Hietanen et al., 2020). Hence, it remains unclear whether the degree of social presence affects social cognition and if these effects are due to the interactivity (here called “immediacy”) or to the amount of information transferred and the shared physical space (called “intimacy”) (Cui et al. 2013; Grondin et al. 2019). Therefore, by examining the impact of social presence on empathy for the first time, we add a potentially important dimension to the study of social cognition. We show that the effects are nuanced: Only the immediate mid-frontal theta response to others’ pain, presumably relating to emotional arousal (Balconi et al. 2009), and SCR coupling were affected by the reduced intimacy. These findings could indicate that intimacy is especially important for more automatic, stimulus-driven empathy components. Future studies should address whether the immediacy within the interaction might have a stronger impact on all components of empathy (Hamilton and Lind 2016; Hietanen et al. 2020).

To conclude, we do not find evidence that empathy for pain is markedly impaired in video-mediated interactions, although physiological and neural resonance with the other’s pain was reduced, which implies that some level of synchronization with the other is impaired. This suggests that empathic abilities might be preserved in everyday mediated social interactions, which are becoming more common. By showing specific changes in empathy components in a mediated interaction, we start to fill the gap in knowledge about social presence in social neuroscience.

### Limitations of the study

The main limitation of the current study is the small sample size, which was due to the complexity of the design. This constraint is especially prominent when reporting mostly null results, as one might argue that our small sample size prevented us from detecting subtle effects of social presence. However, as power analyses showed, by analysing single trials and using a within-subject design, we had sufficient power to detect meaningful effects of social presence. Moreover, the small number of men in the sample prevented us from assessing effects of gender. However, even if there are differences in empathy between female-female and female-male dyads, effects of social presence on it should still be detectable in our within-subjects design. Nevertheless, as the literature suggests possible gender differences in empathy (Christov-Moore et al. 2014), the overrepresentation of women in our sample might prevent a generalization to men. Future studies should therefore assess the effects of social presence on empathy in gender-balanced samples. As we assessed only young students, who were used to video calls in their daily lives, our results might not be generalizable to populations who are less familiar with computer mediated communication. The effects of social presence might be stronger in these populations and future studies should therefore include more diverse samples. Our sample was fairly homogenous in terms of age and sociodemographic background, as we sampled only young psychology students. Homogeneity between participants might influence perceived similarity within a dyad, which might influence empathic accuracy or even its modulation by social presence (Majdandžić et al. 2016; Han 2018). We did not explicitly assess perceived similarity between participants within a dyad and can assume that this varied slightly between dyads, so that we cannot rule out that this slightly influenced our results. However, we also aimed to test the effects of social presence in a generalizable sample, to be sure the effects do not only hold under very specific conditions. Similar to the effects of familiarity between target and observer, future studies will have to address modulatory effects of similarity and other participant characteristics. Another limitation is the comparably low standardization of our laboratory task. Using a task with real live people, we aimed to capture real-life empathy for pain in the best way possible in the EEG laboratory. At the same time, by using a standardized pain-stimulation protocol, we maintained a high degree of standardization compared to studies using unstructured interactive paradigms (i.e., Levy et al., 2017). Our study therefore answers recent calls for more interactive and contextual experimental methods for researching social interaction (Dumas 2011; Przyrembel et al. 2012; Sonkusare et al. 2019).

## Acknowledgements

UMK is supported by the German Science Foundation (grant number KR3691/8-1). We thank Lou Maria Lütjohann, Celina Mävers, Marthe Mieling, Leah Reinicke, Ellyn Sänger and Jasmin Thurley for help with data collection and pre-processing, Charlotte Petereit for help with the figures, and two anonymous reviewers for their helpful comments.

## Author contributions

Conceptualization: P.P. and U.M.K; Methodology: P.P. and U.M.K; Software: P.P; Formal Analysis: P.P.; Investigation: P.P.; Data Curation: P.P.; Writing-Original Draft: P.P; Writing-Review & Editing: P.P., R.W., A.P and U.M.K.; Visualization: P.P.; Supervision: U.M.K.; Project Administration: P.P. and U.M.K; Funding Acquisition: U.M.K

