## supplementary materials for "Effects of social presence on behavioural, neural and physiological aspects of empathy for pain"

### Formula for the (generalised) linear mixed models

### Empathic accuracy

Data range observer rating: 1-101

Data range condition: categorial factor with two levels “direct” vs. “mediated”

Data range intensity: 1-20

Models with all hypothesised fixed effects to find the best random effects structure

Each model is compared to the previous model without the random effect and the random effect is only retained if it improves the model fit.

Model R1: Observer rating~condition*intensity+(1|target/observer)

Model R2 Observer rating~condition*intensity+(intensity|target/observer)

Model R3: Observer rating ~condition*intensity+(condition+intensity|target/observer)

Model R4: Observer rating ~condition*intensity+(condition*intensity|target/observer)

In this case, Model R3 was better than the other models, but Model R4 did not improve the model fit further.

Therefore, the random effects structure of Model R3 was used to evaluate the best fixed effects structure.

Again, each model is compared with every previous model, to test whether the additional fixed effect improves the model fit.

Model 1: Observer rating ~1+(intensity+condition|target/observer)

Model 2: Observer rating ~intensity+(intensity+condition|target/observer)

Model 3: Observer rating ~intensity+condition+(intensity+condition|target/observer)

With t indicating a specific target, o a specific observer, and s a specific trial (stimulus). Int indicated the intenisity in a specific trial, and cond the condition in a specific trial.

In this case, Model 3 was not better than Model 2, and therefore only the effect of intensity was retained in the final model.

Mathematical formula final model:

$\text{Rating}_{tos} = I + I_t + I_o + int_s * b_{int\text{-}t} + cond_s * b_{cond\text{-}t} + int_s * b_{int\text{-}o} + cond_s * b_{cond\text{-}o} + int_s * b_{int}.$

### Unpleasantness

Data range unpleasantness ratings: 1-101

Data range condition: categorial factor with two levels “direct” vs. “mediated”

Data range intensity: 1-20

Model R1: Unpleasantness rating~intensity*condition+(1|target/observer)

Model R2: Unpleasantness rating ~intensity*condition+(intensity|target/observer)

Model R3: Unpleasantness rating ~intensity*condition+(intensity+condition|target/observer)

Model R4: Unpleasantness rating ~intensity*condition+(intensity*condition|target/observer)

In this case, Model R4 was better than the other models.

Therefore, the random effects structure of Model R4 was used to evaluate the best fixed effects structure.

Again, each model is compared with every previous model, to test whether the additional fixed effect improves the model fit.

Model 1: Unpleasantness rating ~1+(intensity*condition|target/observer)

Model 2: Unpleasantness rating ~intensity+(intensity*condition|target/observer)

Model 3: Unpleasantness rating ~intensity+condition+ (intensity*condition|target/observer)

Model 4: Unpleasantness rating ~intensity*condition+ (intensity*condition|target/observer)

In this case, Model 4 was not better than Model 3, and therefore only the effects of intensity and condition wer retained in the final model.

Mathematical formula of the final model:

$\text{Unpleasantness}_{tos} = I + I_t + I_o + int_s * b_{int\text{-}t} + cond_s * b_{cond\text{-}t} + cond_s * int_s * b_{cond\text{-}t*int\text{-}t} + int_s * b_{int\text{-}o} + cond_s * b_{cond\text{-}o} + cond_s * int_s * b_{cond\text{-}o*int\text{-}o} + int_s * b_{int} + cond_s * b_{cond}$.

**Averaged theta responses**

Data range theta response: -1.24 – 2.06

Data range condition: categorial factor with two levels “direct” vs. “mediated”

Data range intensity: 1-20

Model R1: Theta response~intensity*condition+(1|targets/observer)

Model R2: Theta response ~intensity*condition+(condition|targets/observer)

Model R3: Theta response ~intensity*condition+(intensity|targets/observer)

In this case, neither Model R2 nor Model R3 were better than Model R1, and therefore only the random intercepts were retained to evaluate the best fixed effects structure.

Model 1: Theta response ~1+(1|targets/observer)

Model 2: Theta response ~intensity+(1|targets/observer)

Model 3 Theta response ~intensity+condition+(1|targets/observer)

Model 4: Theta response ~intensity*condition+(1|targets/observer)

In this case, Model 4 was better than Model 3, and therefore the interaction between condition and intensity was retained in the model.

Mathematical formula of the final model:

$\text{theta response}_{tos} = I + I_t + I_o + int_s * b_{int} + cond_s * b_{cond} + cond_s * int_s * b_{cond*int}$.

**Condition effects on interbeat-intervals**

Data range interbeat-interval: -257 - 326.02 ms

Data range condition: categorial factor with two levels “direct” vs. “mediated”

Data range intensity: 1-20

Model R1: Interbeat-interval observer~intensity*condition + (1|target/observer)

Model R2: Interbeat-interval observer ~intensity*condition + (intensity|target/observer)

Model R3: Interbeat-interval observer ~intensity*condition + (condition+intensity|target/observer)

Model R4: Interbeat-interval observer ~intensity*condition + (condition*intensity|target/observer)

In this case, Model R4 was not better than the other models, so only the random effects of intensity and condition were retained to evaluate the best random effects structure.

Model 1: Interbeat-interval observer ~1 + (condition+intensity|target/observer)

Model 2: Interbeat-interval observer ~intensity+ (condition+intensity|target/observer)

Model 3: Interbeat-interval observer ~condition+(condition+intensity|target/observer)

In this case, neither Model 2 nor Model 3 were better than Model 1, so no fixed effects were retained in the final model.

Mathematical formula for the final model:

Interbeat-Interval$_{tos}$ = $I + I_t + I_o + int_s * b_{int-t} + cond_s * b_{cond-t} + int_s * b_{int-o} + cond_s * b_{cond-o}.$

**Condition effects on skin conductance responses**

Data range skin conductance observer: 0 -10.16 microSiemens

Data range condition: categorial factor with two levels “direct” vs. “mediated”

Data range intensity: 1-20

Model R1: Skin conductance observer~condition*intensity+(1|target/observer)

Model R2: Skin conductance observer ~condition*intensity+(condition| target/observer)

Model R3: Skin conductance observer ~condition*intensity+(condition+intensity| target/observer)

Model R4: Skin conductance observer ~condition*intensity+(condition*intensity| target/observer)

In this case, Model R4 was not better than Model R3, so only the random effects for condition and intensity were retained in the model to evaluate the best fixed effects structure.

Model 1: Skin conductance observer~1+(condition+intensity|target/observer)

Model 2: Skin conductance observer ~intensity+(condition+intensity| target/observer)

Model 3: Skin conductance observer ~intensity+condition+(condition+intensity|target/observer)

Model 4: Skin conductance observer ~intensity*condition+(condition+intensity|target/observer)

In this case, Model 4 was not better than Model 3, so only the fixed effects for condition and intensity were retained in the final model.

Mathematical formula for the final model:

Skin conductance response$_{tos}$ $= I + I_t + I_o + int_s * b_{int-t} + cond_s * b_{cond-t} + int_s * b_{int-o} + cond_s * b_{cond-o} + int_s * b_{int} + cond_s * b_{cond}$.

**Interbeat-interval coupling**

Data range observer interbeat-interval: -257 - 326.02 ms

Data range condition: categorial factor with two levels “direct” vs. “mediated”

Data range target interbeat-interval: -273.3 - 288.6 ms

Model R1: Interbeat-interval observer ~Interbeat-interval target*condition + (1|target/observer)

Model R2: Interbeat-interval observer ~Interbeat-interval target*condition + (condition|target/observer)

Model R3: Interbeat-interval observer ~Interbeat-interval target*condition + (condition+Interbeat-interval target|target/observer)

In this case, Model R3 was not better than Model R2, therefore only the random effect of condition was retained in the modle to evaluate the best fixed effects structure.

Model 1: Interbeat-interval observer~1 + (condition|target/observer)

Model 2: Interbeat-interval observer ~Interbeat-interval target+ (condition|target/observer)

In this case, Model 2 was not better than Model 1, so no fixed effects were retained in the final model.

Mathematical formula for the final model:

Interbeat-Interval$_{tos}$ $= I + I_t + I_o + cond_s * b_{cond-t} + cond_s * b_{cond-o}$.

**Skin conductance coupling**

Data range observer skin conductance response: 0 -10.16 microSiemens

Data range condition: categorial factor with two levels “direct” vs. “mediated”

Data range target skin conductance response: 0 - 6.16 microSiemens

Model R1: Skin conductance observer ~condition*skin conductance target +(1|target/observer)

Model R2: Skin conductance observer ~condition* skin conductance target +(condition|target/observer)

Model R3: Skin conductance observer ~condition*targ_SCR_sc_add+(condition+ skin conductance target|target/observer)

Model R4: Skin conductance observer ~condition*targ_SCR_sc_add+(condition*skin conductance target|target/observer)

In this case, Model R4 was not better than Model R3, so only the random effects for condition and target skin conductance were retained in the model to evaluate the best fixed effects structure.

Model 1: Skin conductance observer ~1+(condition+ skin conductance target|target/observer)

Model 2: Skin conductance observer ~ skin conductance target +(condition+ skin conductance target|target/observer)

Model 3: Skin conductance observer ~ skin conductance target +condition+(condition+ skin conductance target|target/observer)

Model 4: Skin conductance observer ~ skin conductance target *condition+(condition+ skin conductance target|target/observer)

In this case, Model 4 was the best model.

Mathematical formula for the final model:

$$\text{SCR-Obs}_{tos} = I + I_t + I_o + \mathit{SCR\text{-}Targ}_s * b_{SCR\text{-}Targ\text{-}t} + cond_s * b_{cond\text{-}t} + cond_s * \mathit{SCR\text{-}Targ}_s * b_{cond*SCR\text{-}Targ\text{-}t} + + \mathit{SCR\text{-}Targ}_s * b_{SCR\text{-}Targ\text{-}o} + cond_s * b_{cond\text{-}o} + cond_s * \mathit{SCR\text{-}Targ}_s * b_{cond*SCR\text{-}Targ\text{-}o} + \mathit{SCR\text{-}Targ}_s * b_{SCR\text{-}Targ} + cond_s * b_{cond} + \mathit{SCR\text{-}Targ}_s * cond_s * b_{cond*SCR\text{-}Targ}.$$

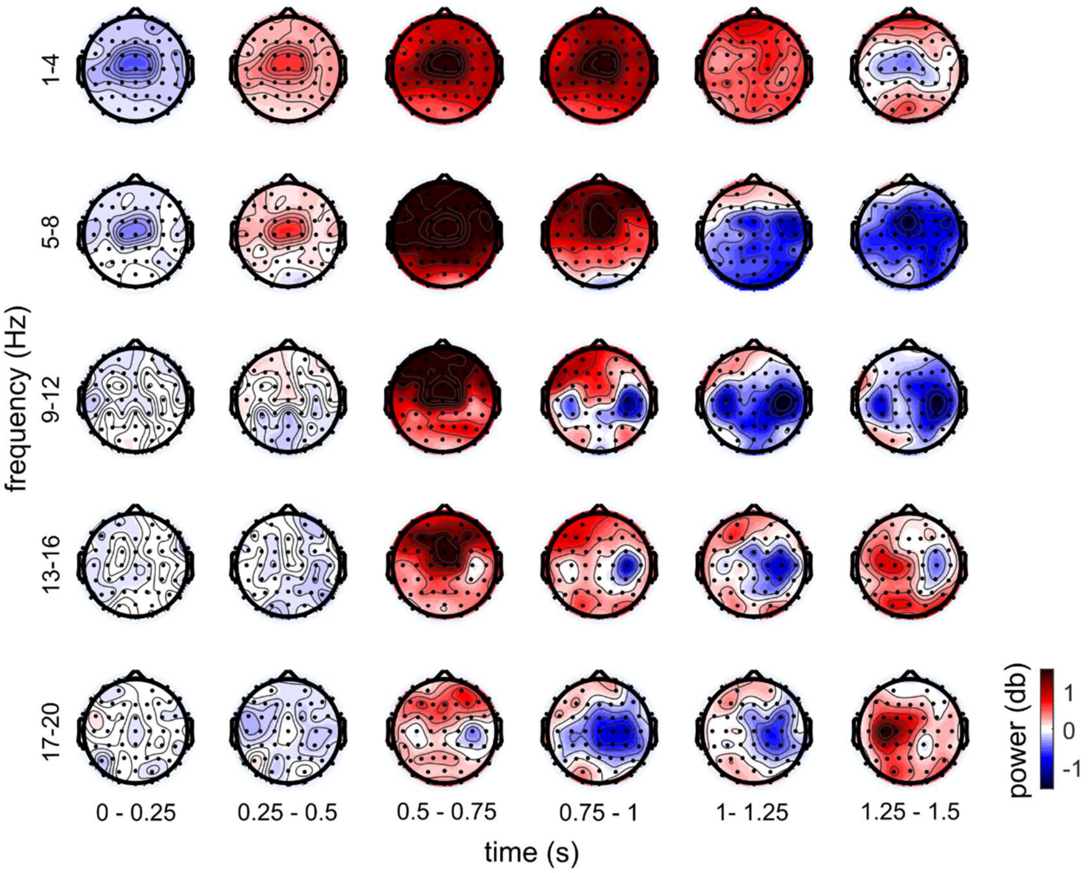


**Figure S1:** Grand averages of the observers' EEG power from 1-20 Hz and from 0-1500 ms after shock onset for low intensity shocks (intensity 1-10) in the own pain condition.

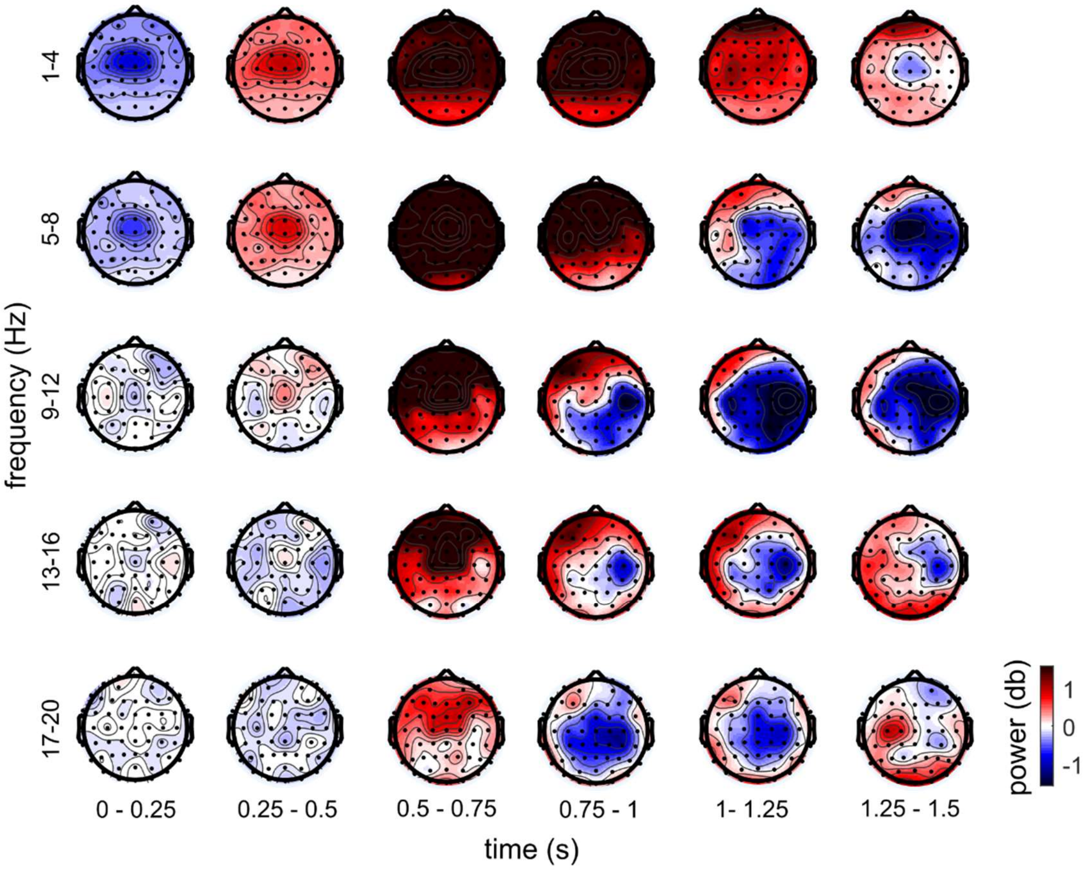


**Figure S2:** Grand averages of the observers' EEG power from 1-20 Hz and from 0-1500 ms after shock onset for high intensity (intensity 11-20) shocks in the own pain condition.

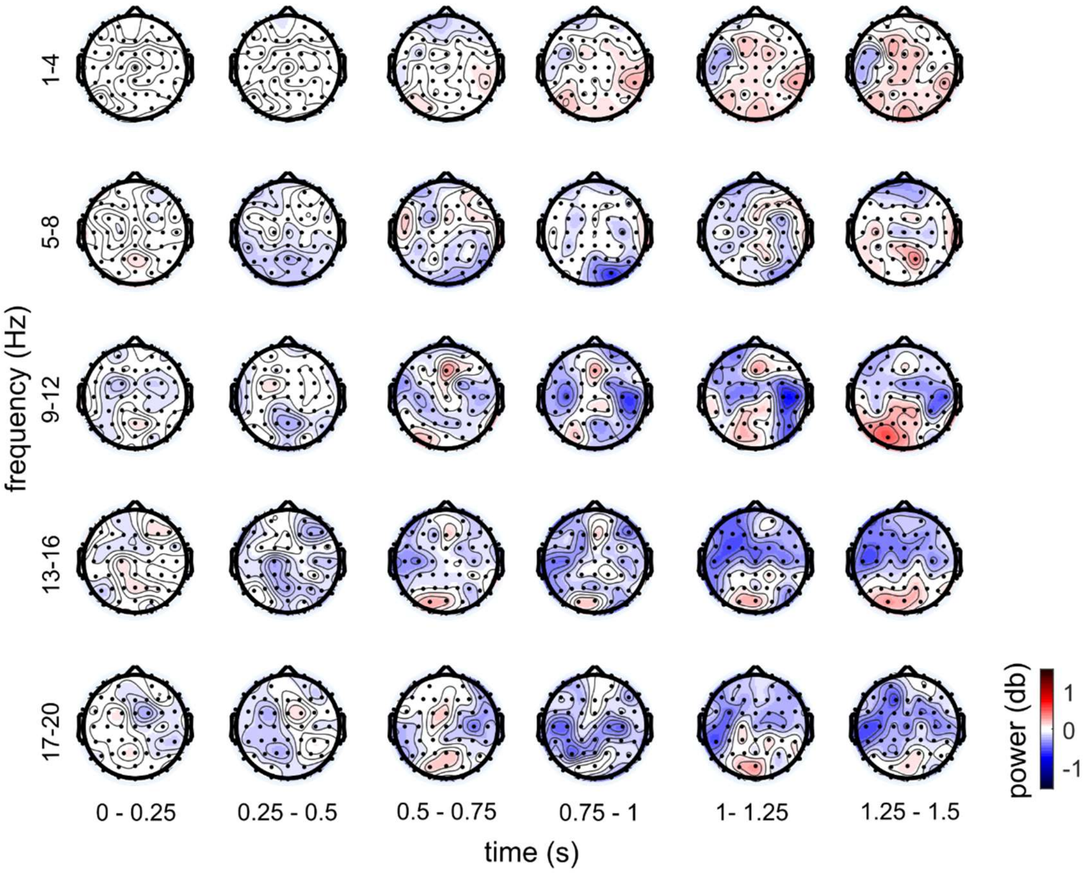


**Figure S3:** Grand averages of the observers' EEG power from 1-20 Hz and from 0-1500 ms after shock onset for low intensity (intensity 1-10) shocks in the direct interaction condition.

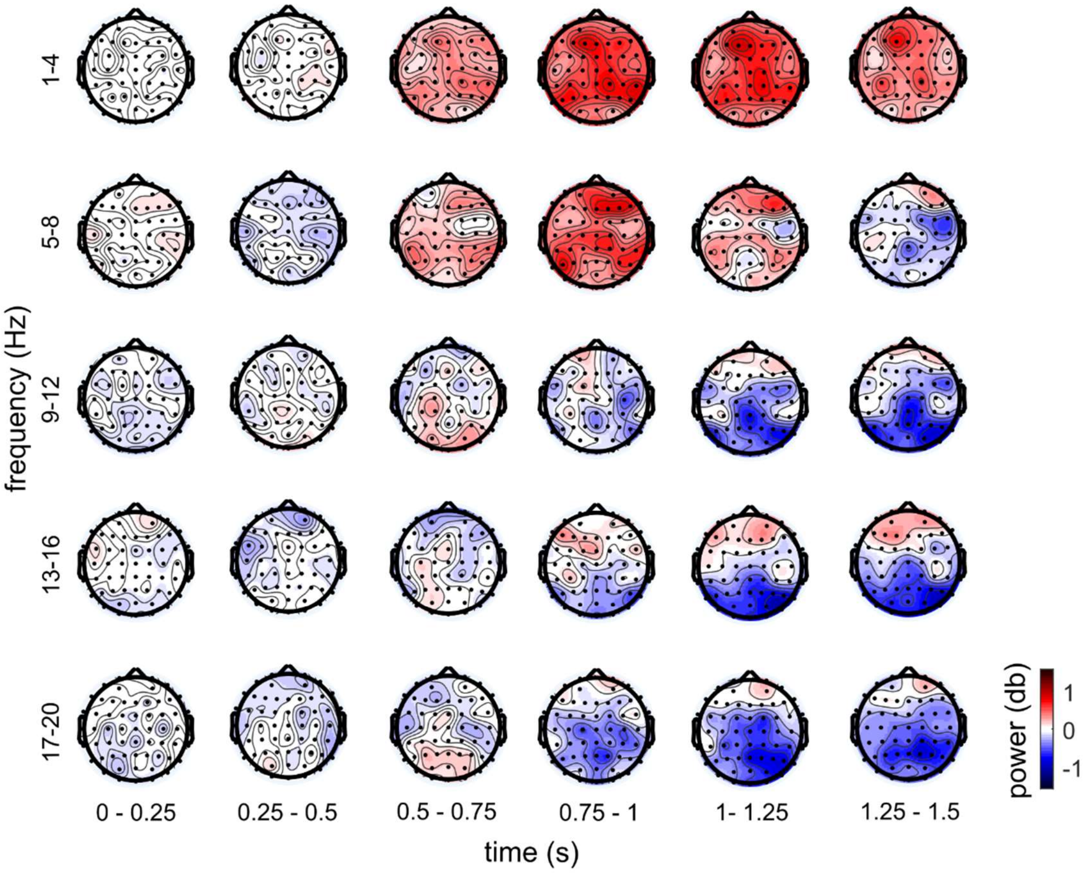


**Figure S4:** Grand averages of the observers' EEG power from 1-20 Hz and from 0-1500 ms after shock onset for high intensity (intensity 11-20) shocks in the direct interaction condition.

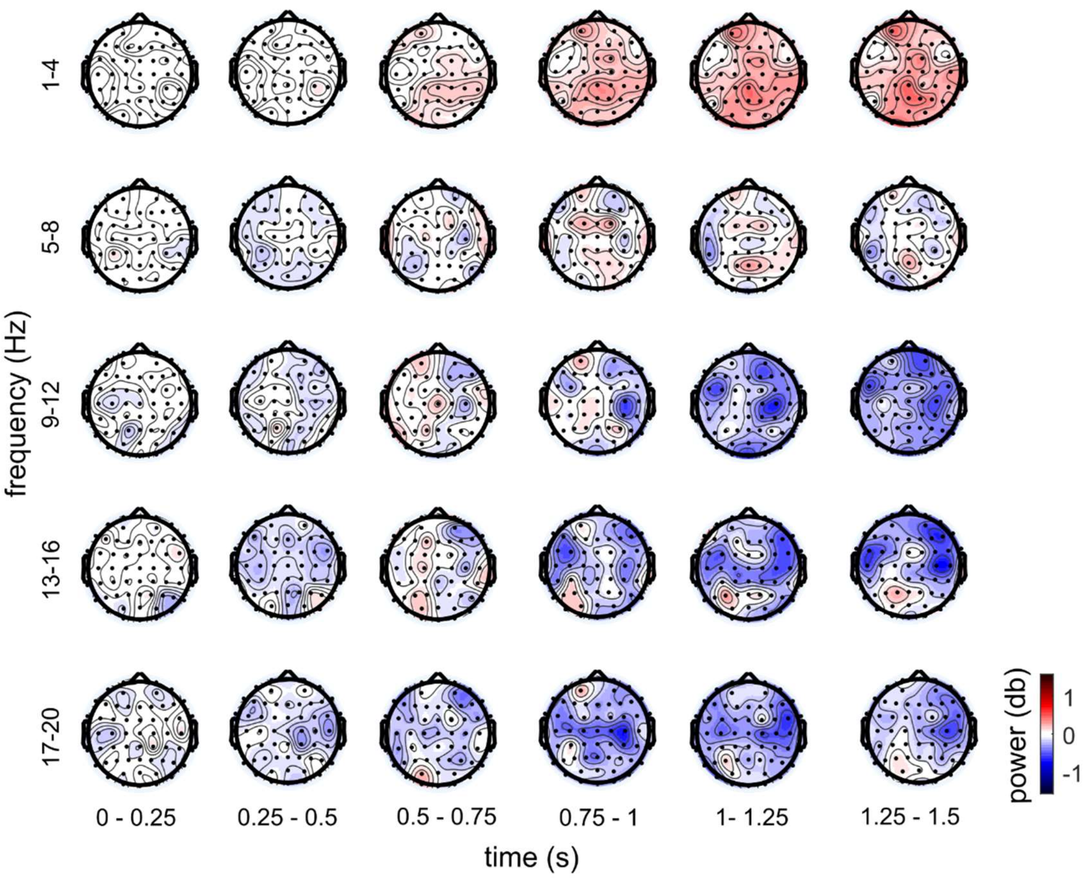


**Figure S5:** Grand averages of the observers' EEG power from 1-20 Hz and from 0-1500 ms after shock onset for low intensity (intensity 1-10) shocks in the mediated interaction condition.

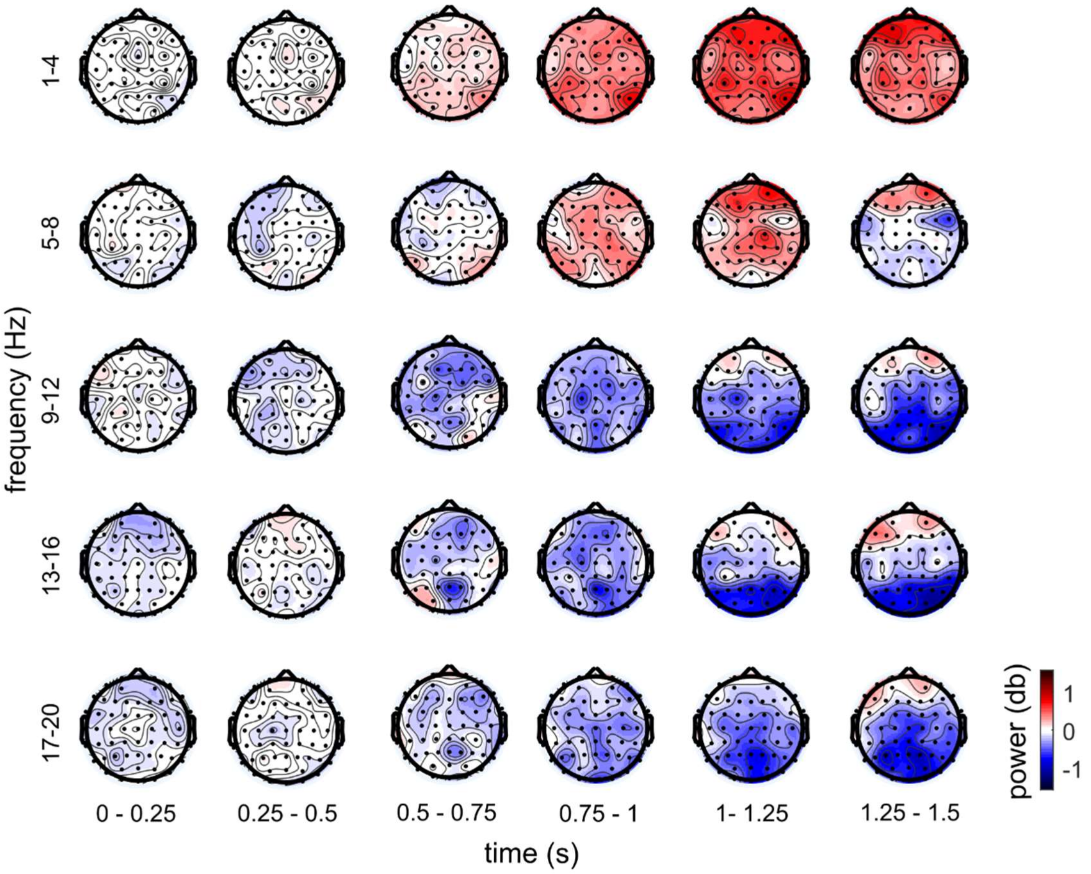


**Figure S6:** Grand averages of the observers' EEG power from 1-20 Hz and from 0-1500 ms after shock onset for high intensity (intensity 11-20) shocks in the mediated interaction condition.

**Supplementary results**

**Familiarity control analysis**

20 of the dyads in our experiment had never met each other before, while 10 dyads were to varying degrees familiar with each other. As past research has shown that familiarity between target and observer can influence empathy, we conducted a control analysis. We hypothesized that in familiar dyads the reduced intimacy in the video-mediated interaction might impact empathic accuracy less compared to unfamiliar dyads, as in familiar dyads observers could also use former knowledge about targets to infer their pain level. We calculated an additional generalized linear mixed model to predict observers' pain ratings from shock intensity and condition, with familiarity and its interaction with the other variables as additional predictors. Familiarity was included as an ordinal variable, as it was derived from participants' answers in the post-experimental questionnaires, which were scaled ordinarily in five steps from "I never met this person" to "I met this person more than 10 times". Although we asked both participants about their familiarity with the other one, we used the familiarity reported by the observer, as the participants did not always agree with each other on this question. We thought the familiarity perceived by the observer would have stronger effects on their experienced empathy.

We report the results of the generalised linear mixed model in the below table.

**Supplementary table 1:** Results of the generalized linear mixed model on observer ratings including familiarity as a predictor

| random slopes | SD | fixed effects | b(SE) | t(df) | p |
|---|---|---|---|---|---|
| intercept/obs | 0.50 | intercept | 2.37 (0.19) | 12.58 | <0.0001 |
| condition /obs | 0.19 | intensity | 0.10 (0.01) | 9.11 | <0.0001 |
| intensity/obs | 0.02 | condition | 0.34 (0.13) | 2.70 | 0.006 |
| intercept/targ | 0.12 | familiarity | -0.89 (0.32) | -2.78 | 0.005 |
| condition/targ | 0.11 | intensity*condition | -0.02 (0.01) | -2.52 | 0.012 |
| intensity/targ | 0.01 | intensity*familiarity | 0.07 (0.02) | 3.77 | <0.001 |
| | | condition*familiarity | 0.42 (0.21) | 2.04 | 0.041 |
| | | intensity*condition*familiarity | -0.03 (0.01) | -1.90 | 0.058 |

The results show that familiarity had a linear main effect on pain ratings, such that observers who had known the targets before judged their pain as lower on average. Moreover, there was an interaction between familiarity and condition, such that the effect of condition on pain ratings was greater for observers who had known the targets before than for observers who had never seen the targets before. There was also an interaction between the effect of intensity on pain ratings and familiarity, indicating that this effect was greater for familiar dyads, suggesting greater empathic accuracy overall in these dyads. However, when fitting follow-up models separately for familiar and unfamiliar dyads, these showed an effect of intensity on observer pain ratings in unfamiliar ($b(SE) = 0.07(0.01)$, $z = 6.24$, $p < .0001$), as well as in familiar dyads ($b(SE) = 0.08(0.02)$, $z = 5.66$, $p < .0001$), indicating meaningful empathic accuracy in the whole sample. In the unfamiliar dyads, there was no main effect of condition ($b(SE) = -0.13(0.10)$, $z = -1.32$, $p = 0.19$) and no interaction between condition and intensity ($b(SE) =-0.003(0.01)$, $z = -0.523$ , $p = 0.60$). Familiar observers judged the others' pain as higher in the online condition than in the presence condition ($b(SE) = 0.32(0.11)$, $z = 2.91$, $p = 0.004$), and were more accurate in the presence than in the online condition ($b(SE) = -0.02(0.01)$, $z = -2.04$, $p = 0.042$). However, as there was a rather small number of familiar dyads, these results should be interpreted with caution. They suggest that familiarity might play a role for the effects of social presence of empathy, but contradict the intuitive hypothesis that video-mediated interaction impacts empathy less if people have already met in real life. Importantly, this suggests that the familiarity between some of the participants did not mask the effects of condition on empathic accuracy, but if anything slightly boosted condition effects. Future studies should either control for familiarity or sample equal numbers of familiar and unfamiliar participants, to explicitly test the effects of familiarity.
